# Unveiling the Hidden Viromes Across the Animal Tree of Life: Insights from a Taxonomic Classification Pipeline Applied to Invertebrates of 31 Metazoan Phyla

**DOI:** 10.1101/2023.11.03.565523

**Authors:** Pau Alfonso, Anamarija Butković, Rosa Fernández, Ana Riesgo, Santiago F. Elena

## Abstract

Invertebrates constitute the majority of animal species on Earth, including most disease-causing agents or vectors, with more diverse viromes when compared to vertebrates. Recent advancements in high-throughput sequencing have significantly expanded our understanding of invertebrate viruses, yet this knowledge remains biased toward a few well-studied animal lineages. In this study, we analyze invertebrate DNA and RNA viromes for 31 phyla using 417 publicly available RNA-Seq datasets from diverse environments in the marine-terrestrial and marine-freshwater gradients. This study aims to (*i*) estimate virome compositions at the family level for the first time across the Animal Tree of Life, including the first exploration of the virome in several phyla, (*ii*) quantify the diversity of invertebrate viromes and characterize the structure of invertebrate-virus interaction networks, and (*iii*) investigate host phylum and habitat influence on virome differences. Results showed that a set of few viral families of eukaryotes, comprising *Retroviridae*, *Flaviviridae* and several families of giant DNA viruses, were ubiquitous and highly abundant. Nevertheless, some differences emerged between phyla, revealing for instance a less diverse virome in Ctenophora compared to the other animal phyla. Compositional analysis of the viromes showed that the host phylum explained over five times more variance in composition than its habitat. Moreover, significant similarities were observed between the viromes of some phylogenetically related phyla, which could highlight the influence of co-evolution in shaping invertebrate viromes.

**Importance:** This study significantly enhances our understanding of the global animal virome by characterizing the viromes of previously unexamined invertebrate lineages from a large number of animal phyla. It showcases the great diversity of viromes within each phylum and investigates the role of habitat shaping animal viral communities. Furthermore, our research identifies dominant virus families in invertebrates and distinguishes phyla with analogous viromes. This study sets the road towards a deeper understanding of the virome across the Animal Tree of Life.

## Introduction

Viruses are arguably the most successful biological entities on Earth, given their ubiquitousness, diversity and numerical abundance, apart from their capability to infect organisms from all domains of cellular life (Breitbart & Rohwer, 2005; Koonin et al., 2006; Wasik & Turner, 2013). In recent times, high-throughput sequencing technologies have catalyzed the identification of new viruses and numerous studies have leveraged metagenomics and meta-transcriptomics to perform discovery of novel viruses at a massive scale (Shi et al., 2016; Porter et al., 2019; Starr et al., 2019; Wolf et al., 2020; Wu et al., 2020; Neri et al., 2022; Zayed et al., 2022). Viruses identified through these approaches enrich reference databases and even are incorporated into the taxonomy of the International Committee on Taxonomy of Viruses (ICTV) (Simmonds et al., 2017a; Simmonds et al., 2023), contributing to a better comprehension and representation of the virosphere.

Thanks to these methods, our understanding of invertebrate viromes, which constitute a vastly more diverse and abundant group when compared to vertebrates, is ongoing unprecedented expansion (Zhang et al., 2018). However, despite the accelerated pace of virus discovery through metagenomics and metatranscriptomics, knowledge of animal viral diversity is still skewed towards viruses that cause disease in humans, involve arthropod vectors, or affect economically relevant species, including culturable arthropods and mollusks (Harvey & Holmes, 2022). Countless viral species, which might play pivotal roles in underexplored organisms, still remain obscured by this bias (Cobbin et al., 2021).

The scientific community is actively making efforts to address this bias. In addition to more conventional studies on disease-vector arthropods (Shi et al., 2019; Shi et al., 2020; Meng et al., 2019; Kondo et al., 2020; Huang et al., 2021), researchers have begun to characterize the viromes of sponges (Pascelli et al., 2020; García-Bonilla et al., 2021), cnidarians (Grasis et al., 2014; Brüwer & Voolstra, 2018; Lewandowska et al., 2020), soil-inhabiting nematodes (Vieira et al., 2022), and echinoderms (Hewson et al., 2020). Some works have greatly improved the exploration of invertebrate viromes with a larger scope, simultaneously analyzing the viromes of invertebrate species distributed across different phyla, including the first characterizations of annelid, platyhelminth and tunicate viromes (Gudenkauf & Hewson, 2016; Shi et al., 2016). However, the Metazoa Tree of Life encompasses a vast diversity of groups (Dunn et al., 2014), and viral communities have yet to be discovered for the first time in many phyla (Harvey & Holmes, 2022).

The SRA archive (https://www.ncbi.nlm.nih.gov/sra) hosts millions of publicly accessible meta-transcriptomic and meta-genomic datasets that comprise more than 7⋅10^16^ sequenced bases (https://www.ncbi.nlm.nih.gov/sra/docs/sragrowth/). Since many researchers who create these datasets do not focus on viral communities, this data provides an opportunity to explore virus diversity in uncharted organisms and ecosystems (Olendraite et al., 2023). The insightfulness of such analyses depends on the quality of complementary metadata in the databases (Pérez-Riverol et al., 2019), including accurate descriptions of sampling conditions, geographic location, and host taxonomy, among other factors. In the present study, the viromes of 417 publicly accessible invertebrate RNA-Seq datasets spanning 31 animal phyla are inferred through a read taxonomic classification pipeline. After classifying the read sequences to existing families of RNA and DNA viruses, the study: (*i*) provides the first description of the virome composition at the family level for several invertebrate phyla, such as Bryozoa, Ctenophora or Onychophora, (*ii*) involves diversity measurements for the estimated viromes and the characterization of the obtained virus-invertebrate interaction network structure, and (*iii*) investigates whether virome composition differences among samples can be attributed to the host phylum and/or habitat.

## Materials and Methods

### Sampling of sequencing datasets

All sequencing accession numbers for the 31 different invertebrate phyla stored in NIH’s SRA database were retrieved in January 2023. This comprehensive database combines information from NCBI’s SRA, EBI’s ENA and DDBJ databases, according to the International Nucleotide Sequence Database Collaboration (INSDC) initiative. This added up to a total of 762,540 accessions for which metadata from multiple fields were fetched from SRA and BioSample databases. Mainly, it was important to collect for each accession its associated BioSample identifier, the organism’s TaxID, the number of sequenced bases, the average read length and details about the library, such as its strategy, source, and layout. Additionally, characteristics of the sampling conditions were retrieved from *isolation_source*, *geo_loc_name* and *lat_lon* fields from BioSample data.

The taxonomic lineage of each TaxID, spanning from the phylum level to subspecies, was obtained using TaxonKit and the NCBI’s taxonomy database (*taxdump*) version last modified on March 2^nd^, 2023. Since NCBI’s species names often involve subspecies, laboratory strains or species mixtures, it was necessary to perform a harmonization step. Namely, all species names had to be binomial, and when this degree of specificity could not be reached, the “sp.” or “spp.” abbreviations would be added after the lowest certain taxon.

SRA metadata information was used to narrow down the selected accession data to a subset that was eligible for the analysis. The following three criteria were used to do this: (*i*) library strategy was restricted to RNA-Seq, (*ii*) average read length had to be greater or equal than 75 bases and (*iii*) the total number of sequenced bases had to be greater than 3 Gb. The rationale behind these two last criteria is to obtain samples with longer reads in order to have higher sensitivity of read-based taxonomic classifiers and to keep samples where the virome composition estimation is as independent from the sequencing depth as possible, respectively.

This led to a total number of 82,145 eligible run accessions. Moreover, by using the three above mentioned criteria a decline was noted in the number of unique species to roughly one fourth of the original number.

To avoid the overrepresentation of individual species, only one run per species was analyzed. Furthermore, since the distribution of unique species is highly asymmetrical between phyla, the maximum number of analyzed species was set to 20 per phylum when possible, trying to select species from different lineages within each phylum. The sampling algorithm tries to maximize the number of samples selected for rare phyla (*i.e*., phyla with a smaller number of datasets), while maintaining a balanced number of samples for phyla with a greater availability of datasets. However, an exception was made for mollusks (39 observations), since they contained a large number of unique species from a wide range of habitats. Taking this into account, the selected species, as well as their final run accession for the analysis, were sampled randomly. When possible, sequenced samples with less than 30 Gb were prioritized over larger datasets in order to speed up the analysis.

The habitat of each organism was reconstructed using the descriptors about the location and source of each sample’s isolate alongside with the information found in the corresponding WoRMS website (Ahyong et al., 2023) entry. Habitat annotation for 143 samples relied solely on ecological information of the species obtained from WoRMS or specific publications, since the geographical location was completely unrecoverable from their metadata in SRA and BioSample. We classified each sample’s habitat into five categories: brackish, freshwater, intertidal zone, marine, and terrestrial. This also served as a way to identify samples that were laboratory-reared, finding that nine out of the 417 analyzed accessions had a laboratory origin. This fraction could be even larger due to the widespread sparse metadata annotation.

### **D**ownload and pre-processing of raw read files

Raw read files were downloaded in FASTQ format using fasterq-dump of SRA toolkit version 2.11.2 (https://github.com/ncbi/sra-tools). A sequential three-step pre-processing phase was conducted over the downloaded raw reads.

First, clumpify.sh script of BBMap package version 38.96 (https://sourceforge.net/projects/bbmap/) was run to remove exact duplicate reads. These can be caused by technical issues of NGS and do not reflect the true abundance of the transcript within the sample’s RNA molecule population. This step was implemented with the purpose of reducing the volume of reads for downstream analyses, which speeds up the pipeline.

Second, SortMeRNA version 4.3.4 (Kopylova et al., 2012) was used to filter out reads of abundant transcripts with non-viral origin. All reads were compared against SortMeRNA’s default rRNA databases for 18S and 28S rRNA subunits of eukaryotes and a custom database consisting of more than 4000 variants of cytochrome c oxidase (*COX*) mRNA sequences. If a single-end read showed significant similarity to any sequence in these reference databases, it was excluded. Similarly, paired-end reads were only kept for downstream analyses if neither of the ends had a match with any sequence in the reference databases.

Third, Trimmomatic version 0.39 (Bolger et al., 2014) was used over the remaining reads to trim out adapters introduced by NGS platforms and terminal regions with an average base call accuracy lower than 80%. The minimum read length after trimming was set to 25.

In some instances, pairing of FASTQ files was found to be corrupted either from the beginning or after any of the preprocessing steps. When necessary, pairing was restored using BBMap’s repair.sh script in order to resume the pipeline.

### Read-based taxonomic classification

The taxonomic classification of preprocessed reads was conducted using Kaiju version 1.8.2 (Menzel et al., 2016). A subset of NCBI’s RefSeq non-redundant proteins (*nr*) database, which includes all protein sequences classified under the superkingdom ‘Viruses’ (TaxID = 10239), served as the reference database. This subset was derived from a version of the *nr* database downloaded in November 2022.

Kaiju takes the reads and translates them to all six possible reading frames. Subsequently, these amino acid sequences are compared with those in the reference database. The best match was stored only if the alignment reached an *E*-value < 10^-5^ threshold.

When a read has a significant viral match, it is added to the count of its particular viral family. However, if the virus of the matching sequence is not classified to a taxon as specific as the family level or if it is not classified under an existing viral family, the read count is added to a category of viruses with uncertain family. So, a vector of read absolute abundances of viral families is constructed for each sample and these are combined into a matrix.

It is important to acknowledge that the virus family nomenclature used in this study is the one reported on the NCBI’s taxonomy database. This database does not incorporate recent updates that have been determined by the ICTV. As a result, it may lack the most recently approved names or, alternatively, include family names that have already been discouraged by the ICTV.

### Statistical analysis of ecological diversity metrics

⍺-diversity at the family level, which refers to the richness of viral families, and Shannon’s diversity index (*H’*) were computed for all estimated viromes restricted to eukaryotic families. ⍺-diversity was calculated as the number of viral families with positive read counts for each sample. *H’* was calculated for each sample *i* as 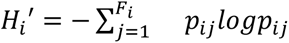, where *F_i_* is the number of families found in sample *i* and *p_ij_* is the proportion of viral reads that are assigned to family *j* in sample *i*.

To find differences between phylum and habitat groups, ANOVA models were fit separately for both of these univariate diversity metrics. To ensure sufficient group sizes, the dataset was reduced by excluding phyla and habitats represented by less than five observations. For both ⍺-diversity and *H’*, the best model was selected with the criterion of minimizing the AIC (Akaike information criterion) after considering all possible combinations of phylum and habitat as covariates.

Since group sizes were unbalanced for both factors, models were checked for homoscedasticity with Levene’s test. In the case of *H’*, phylum grouping was seen to introduce heteroscedasticity in the model, so the data were transformed using the inverse of the Box-Cox transformation of *H’* with parameter λ = 2.5 before the model was fit.

Tukey’s *post-hoc* tests were conducted over the selected models using the phylum as covariate to identify which phyla were responsible for driving the observed differences, if any. Several additional metrics from the field of community ecology were computed after transforming the counts matrix to relative abundances, which indicate the intensity of the ecological interactions. Also, the bipartite network was restricted to the 30 eukaryotic viral families that reached an average relative abundance higher than 0.01% among the samples.

At the network level, measures of (*i*) linkage density, (*ii*) *H*_2_*’* specialization index (Blüthgen et al., 2006), (*iii*) nestedness using the method described by Almeida-Neto et al. (2008), and (*iv*) *Q* modularity index (Newman, 2006) were calculated. One-thousand null models were run to simulate estimates of these four indices under randomness of interactions. The sampling distributions of the simulated indices were used to perform *z*-tests for the observed values. In the case of indices with closed intervals of possible values (*ii* - *iv*), it was checked that the observed value was separated enough from the boundaries to avoid the violation of the normality assumption.

At the level of individual viromes, the *d’* specialization index (Blüthgen et al., 2006) was calculated and used to fit an ANOVA model grouping *d’* estimates by the host’s phylum analogous to the ones described above. *d’* was also calculated for each viral family in the matrix.

### Virome composition analysis

The absolute abundances matrix, which was originally measured in read counts, was first restricted to families of eukaryotic viruses. The vector of abundances of each sample can be regarded as a composition, where each viral family constitutes a part of the virome. In recent years, it has been suggested that microbiome, and by extension virome, datasets should be analyzed under the framework of compositional data analysis (Gloor et al., 2017).

These methods require a processing of the abundance matrix previous to the analysis. In this case, compositions were first transformed to relative abundances vectors with respect to the total number of reads assigned to eukaryotic viral families. In the case of microbiome compositional data, it has also been acknowledged that dimensionality reduction both in observations and features can lead to a more informative analysis (Xia et al., 2018). In this study, it was decided to exclude viral families that did not reach an average relative abundance of 0.01%. Then, imputation of zero values was conducted to avoid the occurrence of null values in the compositions. Finally, the resulting imputed vectors of relative abundances were transformed to centered log-ratio (*clr*) coefficients.

After applying the transformation, the dataset was reduced to the subset of phyla with 15 or more samples. The matrix of Euclidean distances for this subset of *clr*-transformed compositions was used to fit PERMANOVA models (Anderson, 2001). Additionally, PERMDISP (Anderson, 2006) tests were performed alongside PERMANOVA to further investigate the occurrence of centroid location and within-group dispersion differences in the virome compositions between groups. Different PERMANOVA models combining phylum and habitat as covariates were fit and compared in terms of AIC to select the best model.

### PERMANOVA bootstrap analysis of differences between virome compositions

Using the model formula selected after the process detailed in the previous section, a bootstrap implementation of PERMANOVA is proposed to test the existence of differences between the virome composition centroid locations for all pairs of phyla in the dataset.

The use of unbalanced group sizes is known to cause issues when fitting PERMANOVA models (Anderson & Walsh, 2013). However, group sizes are heterogeneous in this study. In an effort to use all available data and keep balanced group sizes, 1500 bootstrap iterations were run using a resampling algorithm with the following steps: (*i*) sample 15 observations per phylum allowing for replacement, (*ii*) impute a new phylum for a random observation (if there are duplicate observations, replace the true phylum in all of them), (*iii*) exclude observations from the imputed phylum until a group size of 15 is reached, making sure the shuffled observation(s) is/are kept, (*iv*) sample new observations for the shuffled observation’s old phylum to restore the balanced group sizes for all phyla (this step is restricted to not sample again the shuffled observation), (*v*) transform the absolute abundance matrix to a matrix of *clr* coefficients, (*vi*) PERMANOVA models and PERMDISP tests are performed for every pairwise combination of phyla, ignoring self and symmetrical comparisons. This results in 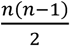 combinations per bootstrap iteration, where *n* is the number of phyla. For each combination, the resulting *P*-values of PERMANOVA and PERMDISP are computed and adjusted by the FDR method considering the total number of tests per iteration. These results are discretized based on the FDR being greater or lower than 0.05. Finally, (*vii*) the combination of acceptance/rejection of the hypotheses of both tests are encoded by a factor of four levels that is stored for each iteration and combination of phyla in an array before proceeding to the next iteration.

### Statistical analysis software

R version 4.0.4 was used for all the statistical analyses downstream of the bioinformatic pipeline. R programming was done using RStudio version 2022.12.0.

In the analysis of species-level ecological measures, the implementation of Levene’s test in car package (version 3.1.2; Fox & Weisberg, 2019) was used to check the assumption of homoscedasticity of ANOVA models. Library multcompView (version 0.1-9; Graves et al., 2023) was used to obtain the compact letter display groups for the results of Tukey tests. Also, library MASS (version 7.3-60; Venables & Ripley, 2002) was used to perform the Box-Cox transformation needed for the *H’* ANOVA model.

The calculation of other measures of community ecology both at network level and at invertebrate virome level was performed entirely using the R library bipartite (version 2.18; Dormann et al., 2009). Null models were simulated using the “r2d” method for quantitative matrices implemented in nulls() function. Both observed and simulated values of the indices were calculated using (*i*) networklevel() with method “linkage density” for the linkage density estimate, (*ii*) H2fun() for the overall specialization index, (*iii*) nested() with method “NODF2” for the nestedness, and (*iv*) computeModules() for the *Q* modularity index. *d’* specialization index was computed using dfun().

In the compositional analysis, libraries zCompositions (version 1.4.0; Palarea-Albaladejo & Martín-Fernández, 2015) and compositions (version 2.0.4; van den Boogaart & Tolosana-Delgado, 2008) were used to prepare the *clr* coefficients matrix with functions cmultRepl() using method “CZM” for the replacement of null values and clr() to perform the *clr* transformation, respectively. Functions adonis() and betadisper() of R library vegan (version 2.6-4; Oksanen et al., 2022) were used to fit PERMANOVA models and perform PERMDISP tests, respectively.

Graphics were generated in R using R base, ggplot2 (version 3.3.5; Wickham, 2016), bipartite, heatmap3 (version 1.1.9; Zhao et al., 2014), and igraph (version 1.2.8; Csardi & Nepusz, 2006) packages.

## Results and discussion

### Characteristics of the RNA-Seq datasets

In the obtained collection of RNA-Seq datasets, the average sample was sequenced for 8.8 Gb and 78% of the samples fell within the range of 3 to 10 Gb. With respect to read lengths, only 22% of the samples had average read lengths outside of the range of 100 to 150 bases. Furthermore, the transcriptomes were obtained using single-end libraries only for 15 samples, while the rest were based on paired-end sequencing.

Pre-processing produced a decline in the total number of reads of at least 45% in all samples. As a result, the number of reads used as the source for taxonomic classification ranged between 5 and 50 million for 86% of the samples (Figure 1A). More detailed information about the pre-processing results is available in the supplementary materials.

**Figure 1.**
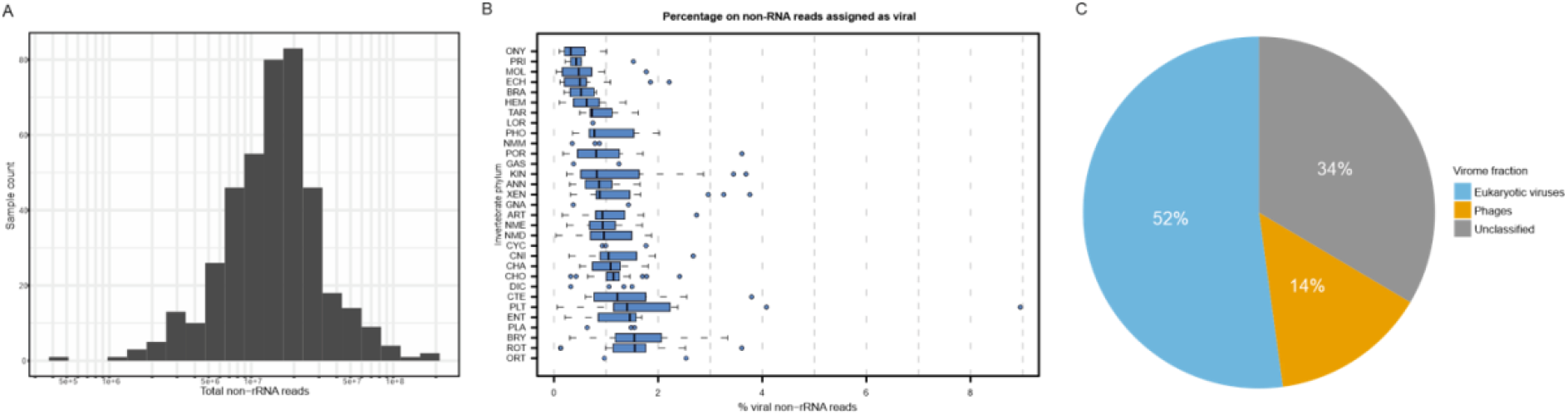
Overview of the bioinformatics pipeline results. **A.** Logarithmic scale plot for the distribution of the total number of reads remaining after pre-processing for each sample. These are the sets of reads that are used as input for the taxonomic classification step. **B.** Distribution of the percentage of reads that showed significant similarity to any viral protein sequence in the *nr* database. Full phylum names can be found in Table S2 at [Supplementary_files]. **C.** Average percentages of viruses by host domain in the whole dataset. Host inferences were made based on the viral family annotation of each hit. The unclassified fraction of the virome comprises viruses of uncertain family.

### Taxonomic distribution of the selected samples

The sampling depth of the different invertebrate phyla exhibited significant variability (Figure 2C). Nine phyla were represented by five or fewer samples, while 16 out of 31 phyla were each represented by at least 15 samples of unique invertebrate species. This variation is due to the differences in the number of species with publicly available transcriptomic datasets (see Table S1 at [Supplementary_files]). Two main factors contribute to this: (*i*) the attention and research focus given to certain species based on anthropocentric interests and (*ii*) the existing biodiversity within each phylum. In fact, it is noteworthy that 83% of all described invertebrate species as of 2013 belonged to the phylum Arthropoda (Zhang, 2013). In the same report, it was stated that species of Arthropoda, Mollusca, Platyhelminthes, Nematoda, Echinodermata, Annelida, Cnidaria, Bryozoa, and Porifera collectively accounted for 98.6% of all invertebrate species.

**Figure 2.**
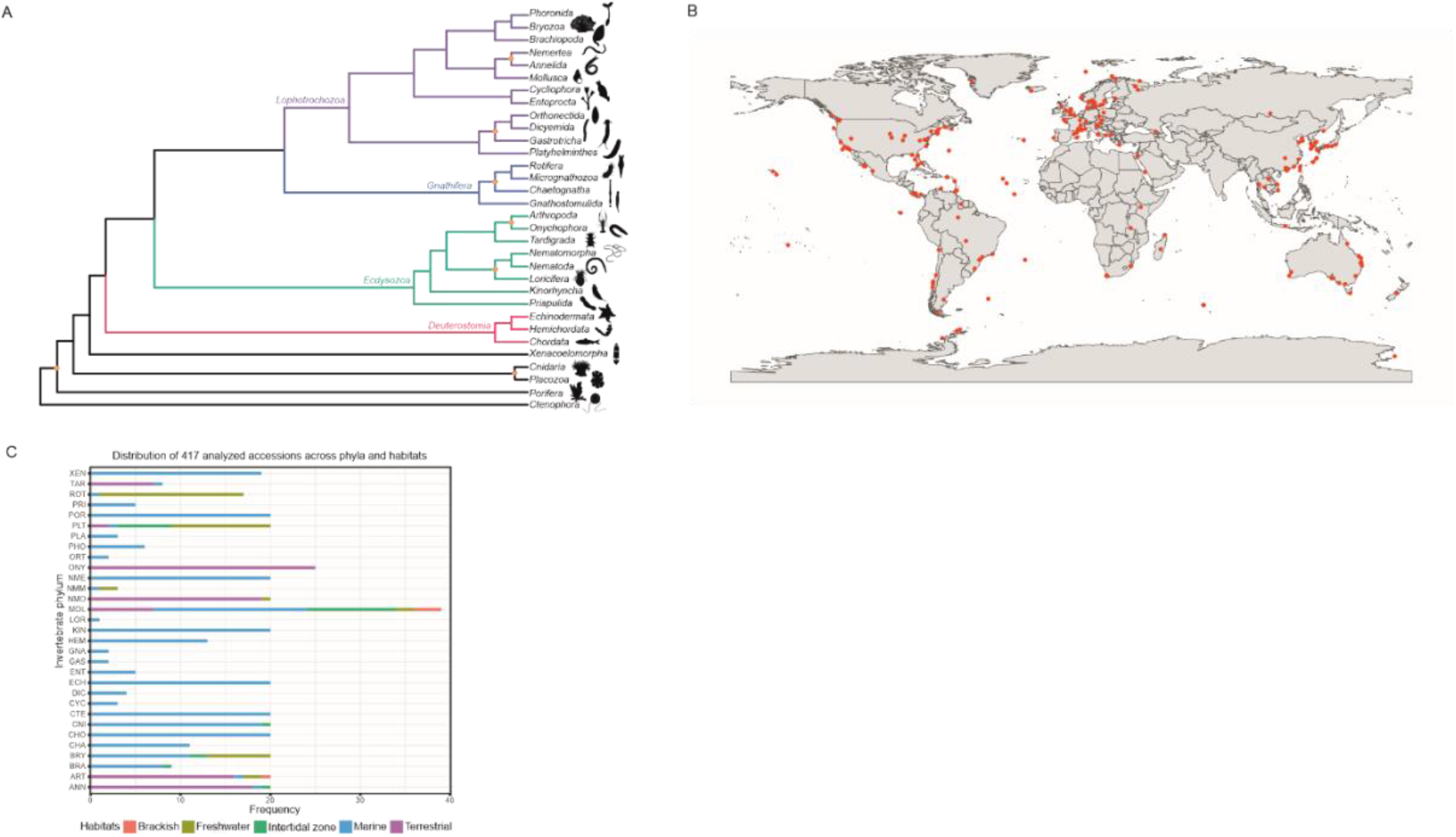
Diversity overview of the invertebrate samples **A.** Approximate topology of the Animal Tree of Life based on our current understanding on animal phylogenetic interrelationships. Polytomies were arbitrarily avoided, opting instead for presentation of nodes with low support or conflicting information across the literature with orange circles. **B.** Geographical locations of the analyzed samples. Latitude and longitude coordinates were originally available for 105 accessions and were approximately reconstructed for further 169 accessions using information from other metadata fields. **C.** Distribution of selected samples across different animal phyla and habitats. Sampling depth was limited to 20 species for all phyla, except for Onychophora and Mollusca. Full phylum names can be found in Table S2 at [Supplementary_files].

A phylogenetic tree for the Animal Tree of Life is presented in Figure 2A to assist in the interpretability of the results. The topology of this tree is mainly based on the results of Laumer et al. (2019). However, in order to include the full set of 31 phyla considered for this study, other works that specifically addressed the placement of certain phyla were taken into account, which was the case for Kinorhyncha, Loricifera and Nematomorpha (Yamasaki et al., 2015) and for Dicyemida and Orthonectida (Lu et al., 2017). Certain nodes that are proposed only in new studies, supported by relatively low bootstrap values or that are in contradiction between different analyses in the literature were highlighted as still disputed/problematic (Schultz et al., 2023; Marlétaz et al., 2019; Juravel et al., 2023; Laumer et al., 2018; Herranz et al., 2022; Wu et al., 2023), since this tree is presented for guidance only.

### Habitat distribution of selected samples

Geographically, the chosen samples covered a wide range of locations around the world, with a notable preference for Europe, North America, Eastern Asia, and Australia (Figure 2B), although it should be noted that the observed spatial distribution across the globe could have been biased by the lack of coordinates for roughly a third of the samples. Besides their ecological habitat, geographic location is acknowledged to shape virome compositions of invertebrates. For instance, a recent study comparing viromes of marine annelids, arthropods and mollusks across different seas found sea-specific virus community profiles (Zhang et al., 2022). Therefore, it is crucial to avoid the over-representation of specific locations before drawing conclusions at a global scale.

With regards to the habitat imputations, sampling depth was also largely variable by habitat (Figure 2C). The marine habitat constituted the most abundant category with 257 samples, followed by the terrestrial habitat with 94 observations. The freshwater habitat was less common, with 41 samples, while the intertidal zone and brackish habitats were even scarcer, with 21 and 4 samples, respectively. For 20 out of the 31 analyzed phyla, there was only one observed habitat, and in most of these cases, it was the marine habitat, except for Onychophora, where all samples were terrestrial. Habitat variability was remarkable in Arthropoda, Bryozoa, Mollusca, and Platyhelminthes.

### Read taxonomic classification into viral families

Overall, the proportion of the pre-processed subset of reads with significant similarity to any viral protein sequence was lower than 2% for approximately 92% of samples (Figure 1B). On average, 1% of reads were assigned as viral. There is an outlying sample in terms of the viral proportion of the reads population, which corresponds to the flatworm *Rhynchomesostoma* sp. (9%). This was likely influenced by the fact that this was the only sample with less than a million reads, leading to a more inaccurate estimation of its transcriptome. When sorting the phyla based on their median viral proportion of reads, it can be observed that phyla such as Onychophora, Mollusca, and Echinodermata lie on the lower end of viral abundance, while samples of Rotifera and Bryozoa harbor a larger proportion of viral RNA.

Additionally, when examined by habitat, freshwater samples showed the highest proportion of viral reads on average (1.7%), while terrestrial samples had the lowest (0.8%). These results reflect two sides of the same coin, as the higher viral abundance in freshwater samples aligns with the dominance of bryozoans, platyhelminths and rotifers seen above. In contrast, Onychophora, the phylum with the lowest viral abundance, is strictly terrestrial.

The proportion of viral RNA was found to be much more variable in another study focusing on the RNA viromes of several invertebrate phyla, ranging from 0.05% to 87% (Shi et al., 2016). However, in another study on the virome of insect species *Aedes albopictus*, it was determined that the fraction of eukaryotic viral reads, this time including rRNA reads, ranged from 0.2 to 1.8% (Shi et al., 2020). Another publication showed that the viral proportion of reads between samples of two other mosquito species was different both on average and in the range of observed values (Shi et al., 2019). A virus enrichment step was not included in the extraction protocol of any of these studies. Comparatively, the figures reported in the present study are lower and display less variation than what was reported in the literature. However, it is important to note that all of the mentioned studies used taxonomic annotation of contigs instead of reads, which may have had an effect on the results (Tamames et al., 2019).

The majority of reads assigned to viral sequences were classified under families of eukaryotic viruses (52%), while phages (including here viruses of archaea) constituted a minority of the viral RNA in the analyzed samples, making up an average of 14% (Figure 1C). A remarkable proportion of viral reads mapped to sequences of undetermined viruses or virus species that are not yet classified to viral families in the NCBI taxonomy database. This fraction of unclassified viruses ranged between 1% (*Pocillopora damicornis*, Cnidaria) to 69% (*Cladolabes schmeltzii*, Echinodermata) of the whole virome. Among the deeper explored phyla, half of the samples of Ctenophora and Porifera have estimated viromes with less than 25% of unclassified viruses, while for Rotifera and Brachiopoda, half of their samples harbor viromes with approximately 40% or more relative abundance of viruses of uncertain family (see Figure S1A at [Supplementary_files]). When considering habitats, the median proportion of unclassified viruses is 32% for terrestrial and marine habitats, while it reached 38% for intertidal and freshwater samples.

Excluding the unclassified fraction of the virome, the phage proportion was highly variable, even within phyla and habitats: the lowest value was observed for the kynoryhinc *Echinoderes dujardinii* (0.4%), while the sample with the highest relative abundance of phage reads was a sea cucumber, *C. schmeltzii* (99.4%). However, the median phage proportion of the classified virome ranged between 11 to 12% for all three most sampled habitats (freshwater, marine and terrestrial). For most phyla, the phage proportion of the virome showed a positively skewed distribution. The phylum with the highest proportion of phages was Porifera (43.3% on average), which goes in line with recent discoveries on the dominance of viruses of the order *Caudovirales* among the viral communities of sponges (Pascelli et al., 2020).

Out of the 134 viral families identified in the 417 samples, 90 were known to infect eukaryotic organisms. 28 of these families were found only in four or less samples, which represents less than 1% of the whole dataset (see Table S3 at [Supplementary_files]). In contrast, 13 viral families were identified in more than 95% of the samples, and the presence of reads classified to 5 of these families was consistent across all 417 samples, namely *Flaviviridae*, *Marseilleviridae*, *Mimiviridae*, *Phycodnaviridae*, and *Retroviridae*.

The average relative abundance was very uneven among families: *Retroviridae* dominated the eukaryotic viromes, accounting for 34.7% of reads assigned to eukaryotic viruses, followed by *Mimiviridae* at 23.5% (see Figure S1B and Table S4 at [Supplementary_files]). Besides *Retroviridae*, the group of highly abundant viral families was dominated by nucleocytoplasmic large DNA viruses (NCLDV): altogether, *Iridoviridae*, *Marseilleviridae, Mimiviridae*, *Phycodnaviridae*, *Pithoviridae*, and *Poxviridae* constituted on average 41% of reads among the estimated eukaryotic viromes. However, proteins identified within these ubiquitous families often exhibited eukaryotic orthologs or xenologs. For instance, actin-like products were erroneously classified to *Retroviridae* in great numbers (CAA25063.1, P00544.1, QDH76138.1, and QDH76139.1). Furthermore, numerous sequences assigned to NCLDV families exhibited significant similarity to various host proteins, including heat-shock proteins, tRNA synthetase, histone, and ubiquitin sequences, among others. Although these genes with eukaryotic homologs can potentially be encoded by NCLDV (Raoult et al., 2004; Boyer et al., 2009; de Souza et al., 2021; Ha et al, 2021; Farzad et al., 2022; Talbert et al., 2022), their abundance was significantly greater than those associated with the virus replication, which lack close homologs in the host genomes (Koonin & Yutin, 2010). This could hint in the direction of these hits representing host sequences in the samples, which could partly explain why a large part of the apparently more dominant viral families identified across the viromes were DNA viruses, contrary to the common consensus, which states that RNA viruses dominate invertebrate viromes (Shi et al., 2016; Porter et al., 2019).

*Flaviviridae*, an RNA virus family known to infect invertebrates (Moureau et al., 2015; Parry & Asgari, 2019), emerged as the third family with a higher average relative abundance of reads. However, its hits were often linked to a polyprotein of a bovine pestivirus species, incorporating a ubiquitin-like sequence (AAA42855.1). Flaviviruses are known to interact with host’s ubiquitin (Wang et al., 2016) and some flaviviruses of vertebrates even incorporate it into their genomes (Mayer et al., 2003; Simmonds et al., 2017b). However, the scarcity of other matches for *Flaviviridae* and the recent incorporation of ubiquitin to genomes of flaviviruses make it unlikely that a significant number of these transcripts have a true viral origin. Consequently, the abundance of flaviviruses may have been notably overestimated within the viromes.

The surprising position of *Mitoviridae* as the fifth most consistently abundant viral family was found to be almost entirely caused by the identification of a highly conserved transcription elongation factor of eukaryotes that was misannotated as part of a mitovirus genome in the reference database (QDH87640.1).

Overall, it was challenging to determine whether the identified transcripts stemmed from genomes of independent viruses or from integrated virus sequences within the host’s genome since the study was based solely on RNA-Seq data. It is well-known that endogenous viral elements (EVEs) of DNA and non-retroviral RNA viruses are widespread across invertebrate genomes and these EVEs can be transcriptionally active (ter Horst et al., 2019; Veglia et al., 2023). Furthermore, EVEs of invertebrate genomes originate from a wider range of viral families when compared to vertebrate genomes and the abundance and diversity of EVEs is highly variable even between closely related invertebrate species (Gilbert & Belliardo, 2022). Besides, in case of a true presence of retroviruses in the samples, it remains unknown if the viruses are found in the form of proviruses, as it has been reported to be common in insect retrovirus infections (Terzian et al., 2001). It is also uncertain to which extent these integrated viruses could be transcriptionally active across the different invertebrate phyla.

With respect to the host of the detected viruses, 18 of the 30 most abundant viral families (see Table S4 at [Supplementary_files]) have at least one member described that is able to infect invertebrates, considering the recent discoveries in *Hepeviridae* (Wu et al., 2018), *Mitoviridae* (Jacquat et al., 2022) and *Partitiviridae* (Cross et al., 2020). In many cases, the known host range of these families among invertebrates is limited to arthropods, but it is yet unknown if eventually they will be found in other invertebrate phyla. For instance, the presence of viruses of the family *Iridoviridae* was recently confirmed in sponges, thereby broadening its previously acknowledged host range (Canuti et al., 2022). In our results, around 0.8% of all eukaryotic virus reads were mapped to *Iridoviridae* for Porifera samples on average, which is nonetheless lower than the mean relative abundance reported overall (see Table S4 at [Supplementary_files]).

Sequences related to *Herpesviridae* were also identified in substantial amounts, following a trend seen in other studies on invertebrate viromes (Grasis et al., 2014; Feng et al., 2017; Brüwer & Voolstra, 2018). In this case, most of the hits for *Herpesviridae* were mainly related to sequences of thymidylate synthase, which is encoded both by the genomes of herpesviruses and eukaryotes (Gáspár et al., 2002). To a lesser extent, hits to both subunits of ribonucleotide reductase were also frequent for *Herpesviridae*. This enzyme is also encoded by herpesviruses and their hosts and has a key role at different stages of the virus’ replication (Wnuk & Robins, 2006; Dufour et al., 2011). Although *Herpesviridae* is acknowledged to harbor exclusively vertebrate-infecting viruses, it has recently been suggested that the closely related family of *Malacoherpesviridae*, whose only two members are mollusk-infecting viruses, could be largely underexplored (Rosani et al., 2023). In our study, sequences classified to *Malacoherpesviridae* were rare, absent from most of the samples and mostly related to a single hit, a putative ATP-dependent DNA ligase (YP_006908759). This protein is not encoded by viruses of *Herpesviridae*, but it has been annotated with this function in three different malacoherpesvirus genomes. Apparently, this protein shows especially close similarity to the ATP-dependent DNA ligase of invertebrates, mainly arthropods, nematodes, placozoans, and platyhelminths. BLASTp searches were performed for the query sequences that matched this protein and significant similarities to the homologs within invertebrate genomes were found. Thus, it is unclear to which extent the abundance and ubiquitousness of herpes-like viruses was overestimated.

Establishing the hosts of viruses discovered through meta-transcriptomics presents significant challenges, given the coexistence of multiple potential hosts within a single sample (Cobbin et al., 2021). Generally, the microbiome of an invertebrate constitutes a diverse set of organisms that can serve as hosts for very different viruses. Although the bacterial part of the microbiome can be ignored in this context, since phages have been excluded from the analysis, the eukaryotic fraction is nowhere near negligible in invertebrate microbiomes (Holt et al., 2022). The common occurrence of these symbioses could have further contributed to the reported abundance of protist-infecting NCLDVs. Additionally, RNA from eukaryotic viruses can be present in dietary components, ranging from other invertebrates to protists, algae, or plants or they can be present simply due to sample contamination. Host determination lies beyond the scope of the current study, and host-virus associations are only tentatively inferred from the host ranges documented in the existing literature for each family.

The individual virome compositions for all the analyzed samples, as well as the aggregated viromes by phylum and habitat can be accessed in an online resource (provisionally accessible at http://invertviromes.i2sysbio.uv.es/).

### Virome diversity and community structure analysis

On average, approximately 22 different eukaryotic viral families were found for each sample with an interquartile range spanning between 19 and 24. The most extreme cases were the sample of *Botryllus schlosseri* (Tunicata) with 37 identified families and the samples of *Echinoderes dujardinii* (Kinorhyncha) and *Xenoturbella bocki* (Xenacoelomorpha) with only 11 families (Figure 3A). Overall, ⍺-diversity was similar between the samples, a trend congruent with findings from other studies on invertebrate viromes that employed a comparable RNA-Seq analysis pipeline (Brüwer & Voolstra, 2018).

**Figure 3.**
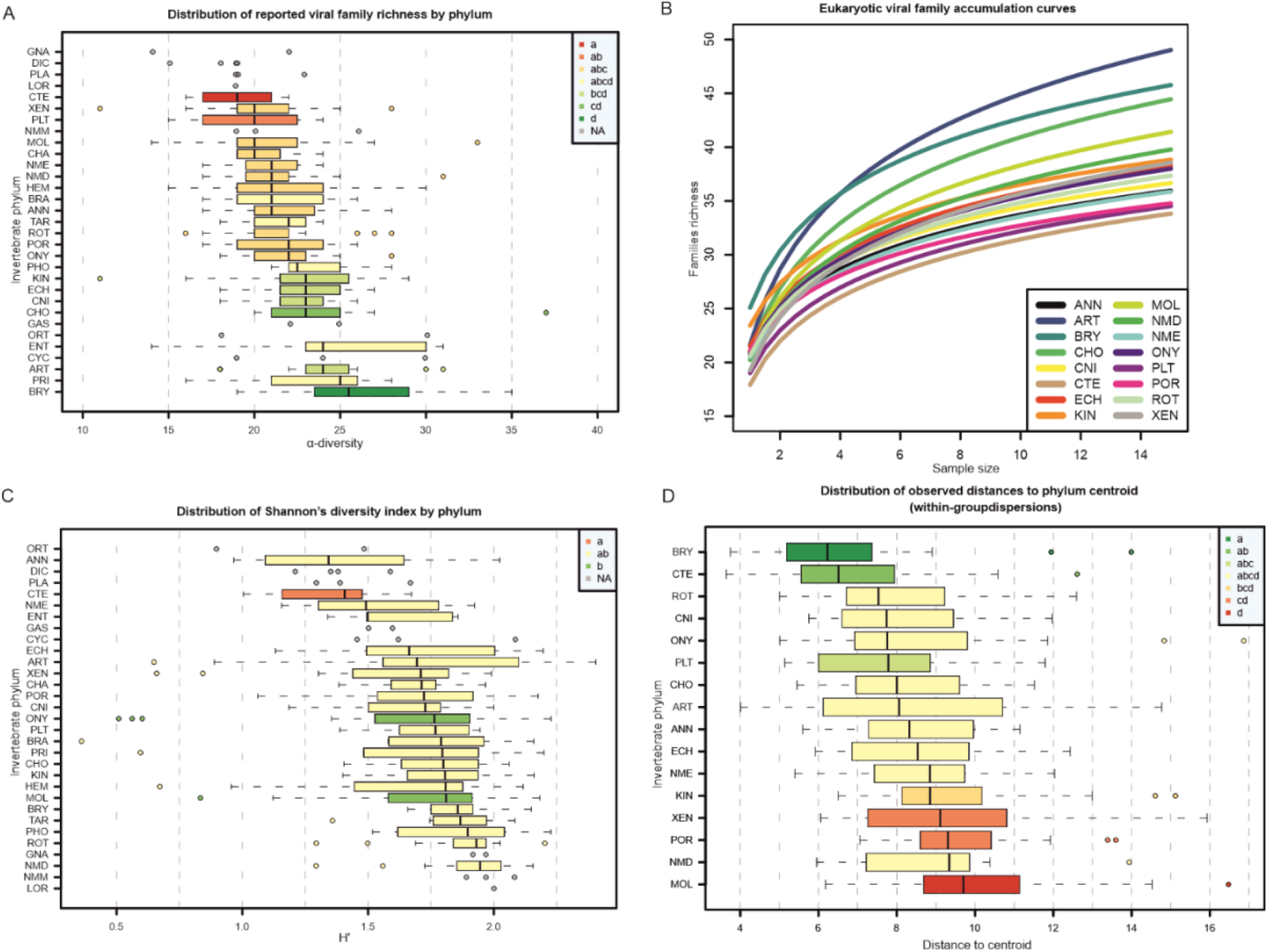
Comparison of the estimated virome diversity between phyla. **A.** Observed distribution of the richness in eukaryotic viral families (⍺-diversity) for the samples of each phylum. Phyla represented by less than 5 samples were excluded from the ANOVA analysis comparing ⍺-diversity between phyla. The legend displays the groups defined by the *post hoc* Tukey test. **B.** Eukaryotic viral families accumulation curves by phylum. At each sample size, 50 iterations were performed to extract different samples of each size and compute their ⍺- diversity. A linear model was fit for the calculated values against the logarithm of the sample size to display the curves. Only phyla with 15 or more samples were included. **C.** Observed distribution of the Shannon’s diversity index for the eukaryotic fraction of the viromes. Phyla represented by less than five samples were excluded from the ANOVA analysis. A transformation of the family Box-Cox, a monotonic increasing function, was performed to avoid heteroscedasticity in the ANOVA model. The legend displays the groups defined by the *post hoc* Tukey test using the transformed values. **D.** Distribution of the within-phylum variability of the virome compositions. The Euclidean distances of the *clr-*transformed compositions were obtained to compute the phylum centroid and the distances of each sample to its phylum centroid. Distances to phylum centroid were analyzed with an ANOVA model for the phyla with 15 or more samples. The legend displays the groups defined by the post hoc Tukey test. Full phylum names can be found in Table S2 at [Supplementary_files].

The ANOVA model for family richness that involved both phylum and habitat as additive factors showed the best fit in terms of AIC. Notably, the influence of phylum on ⍺-diversity differences was significantly more pronounced (*P* < 0.0001) compared to habitat (*P* ≈ 0.05). The significance of regression parameters for habitat groups became lower when phylum was added to the model, which supports the idea that a remarkable degree of collinearity exists between both factors, with phylum grouping being more informative. According to *R*^2^, 23% of the variance in family richness was explained by phylum and habitat groupings together. Homoscedasticity was checked for both factors (*P* > 0.05).

The rejection of the null hypothesis in the ANOVA model suggests that some differences in the average family richness exist between the phyla. *Post hoc* pairwise comparisons revealed that bryozoans showed a significantly higher family richness than the majority of tested phyla (Figure 3A). In sharp contrast, Ctenophora was the phylum with the lowest number of different viral families per sample (Figure 3A). The outlying sample of Chordata (Tunicata) shifted its mean to positive values compared to its median, which makes it the phylum with the second highest number of viral families on average. When seen by habitat, terrestrial and marine samples had an average ⍺-diversity almost identical to the global mean. The average viral family richness for freshwater samples was slightly higher at 23 families and intertidal zone samples showed the lowest ⍺-diversity overall with 21 distinct families on average per virome. Compared to purely marine habitats, intertidal zone habitats pose a series of stresses to local organisms (Mouritsen & Poulin, 2002), which might reduce the range of hosts that viruses are adapted to and can find in intertidal zones. This could help to explain the observed small difference in ⍺-diversity observed between habitats.

Family richness was also analyzed by phylum and habitat using rarefaction curves (Figures 3B and S2A at [Supplementary_files]). This revealed that a number of samples > 15 seems to be required to capture the full range of distinct viral families within any phylum, as none of the curves reached a plateau at this sample size. However, the lack of available samples for some phyla in the SRA database makes it impossible to reach that plateau. This should point out that more sequencing and explorative efforts are needed to describe the full diversity of invertebrate viruses.

In the case of Arthropoda, the accumulation curve seemed to be exceptionally steep, which suggests that the sampled transcriptomes for this phylum harbored the most diverse sets of viral families (Figure 3B). Other phyla that stand out for a higher accumulation of viral families were Chordata (Tunicata), Mollusca and Bryozoa. However, Bryozoa differed from the rest of the cited phyla by starting at a much higher value of ⍺-diversity. On the lower end, Ctenophora, Platyhelminthes and Porifera showed the lowest accumulation of viral families as the sample size increased (Figure 3B). It does not seem that habitat affiliations are the main cause of these differences between phyla, since single habitat phyla were found on both extremes, just as phyla with more variable habitats. Interestingly, two of the phyla with the lowest rate of viral families accumulation are precisely the earliest branching animal phyla: Porifera and Ctenophora (Li et al., 2021).

The observed Shannon’s *H’* values ranged between 0.4 and 2.4, lying in most cases between 1.5 and 2 (Figure 3C). *H’* was more variable than family richness, which suggests that evenness introduced a substantial degree of variation between the estimated viromes. Roughly 4% of the samples had viromes with *H’* < 1, which might have been caused by one or few viral families dominating the virome. This unevenness could be the consequence of ongoing infection, where transcripts associated with infecting viruses become much more abundant.

In this case, the best ANOVA model fit was the one-way model with phylum as the single covariate, which suggests that habitat was not as informative for the observed differences in *H’*. Since Levene’s test showed rejection of the hypothesis of homogeneity of variances across the phyla, a Box-Cox transformation was applied. Although the applied transformation could hinder the interpretation of the Tukey’s *post hoc* test, the use of the untransformed *H’* was ruled out to avoid misleading results. Onychophora and Mollusca appeared to have significantly a higher mean in *H’* when compared to Ctenophora (Figure 3C). But Onychophora and Mollusca did not show exceptionally high *H’* values, it was their group sizes that were exceptionally large. Moreover, Ctenophora appears once again to have less diverse viromes also according to this measure (Figure 3C).

All four network-level measures computed for the bipartite network of viral families and invertebrate species were found to significantly differ from the estimates obtained from the null model (see Table S5 at [Supplementary_files]). In particular, the observed linkage density, nestedness, and *Q* modularity were significantly higher than expected by chance, while the network-level specialization index *H*_2_*’* was lower. In all cases, especially for *H*_2_*’* and *Q*, the rejection of *z*-tests appears to be driven more by the high dimensionality of the matrix rather than substantial differences between observed measures and values estimated under the null model.

The high degree of nestedness in the network might suggest the coexistence of both generalist and specialist organisms on both sides of the matrix. This combination of high nestedness and low modularity is a recurring pattern observed in phage-bacteria and virus-plant networks (Weitz et al., 2013; Valverde et al., 2020). As shown in Figure 4A, the representation of a bipartite matrix by phylum visually highlights nestedness, mainly through the aforementioned group of ubiquitous viral families in contrast to rarer families. This set of ubiquitous families, comprising *Baculo-*, *Flavi-*, *Partiti-*, *Polydna-*, *Retroviridae*, and several NCLDV families, could form a core virome that is common to the majority of the analyzed samples. Nevertheless, nestedness could have been overestimated because of the erroneous classification of highly conserved host sequences as viral, as seen in some of these families.

**Figure 4.**
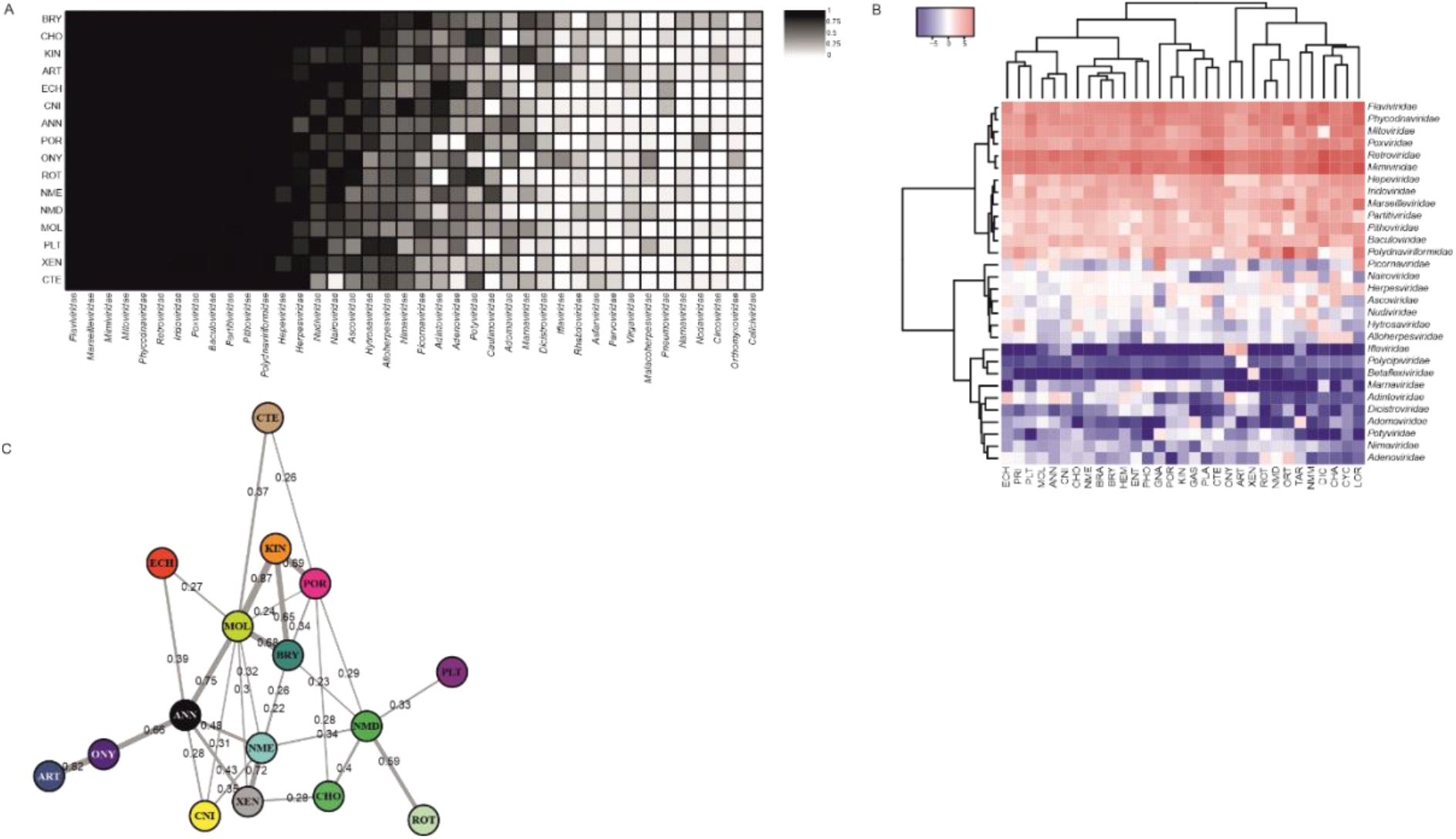
Compositional analysis of the viromes for phyla with 15 or more samples. **A.** Virus- invertebrate interaction matrix by phylum. The color gradient illustrates the proportion of samples by phylum where each particular viral family was present. All 40 viral families shown are present in 20% of the samples of at least one phylum. Ordination of rows and columns highlights the nested nature of the interaction network. Full phylum names can be found in Table S2 at [Supplementary_files]. **B.** Heatmap and hierarchical clustering analysis by phylum of the 30 most abundant eukaryotic viral families. Relative abundance vectors of viral families were aggregated by phylum giving the same weight to each sample prior to performing the *clr*- transformation. The hierarchical clustering of both phyla and viral families was performed using the Ward2 method. The legend of the heatmap displays the scale of the *clr* values. **C.** Virome centroid equivalence network by phylum. Pairwise PERMANOVA and PERMDISP tests were computed in 1500 bootstrap iterations for the *clr* coefficients vectors of each combination of phyla. Differences between phyla were considered significant only when PERMANOVA’s H_0_ was rejected but PERMDISP’s H_0_ was accepted. Links are shown connecting nodes only when the proportion of iterations where differences between phyla were not significant was higher than 0.2. Bootstrap support values are shown for each link.

Specialization indices (*d’*) were calculated for both invertebrate species and viral families within the bipartite matrix. Remarkably, the distribution of *d’* values was almost identical for both viruses and hosts, both having a mean *d’* = 0.01. The highest observed *d’* values did not surpass 0.06 in any case. These low *d’* values suggest that interactions are not restricted to specific pairs of organisms. Consequently, any of the 30 viral families included in this matrix could potentially be found in any of the viromes, as determined by the taxonomic classification pipeline employed. Furthermore, a comparison of d’ indices among phyla using an ANOVA model revealed a significant association between phylum grouping and variations in *d’* values (*P* < 0.0001). Notably, species belonging to Onychophora, Tardigrada, Arthropoda, and Mollusca displayed a higher degree of specialization in hosting viral families. In contrast, Chaetognatha, Nematoda, and Bryozoa were characterized by a greater tendency toward generalism of viral families.

### Virome composition analysis

In order to assess the dominance of each viral family within the viromes, the matrix of relative abundances was transformed into *clr* coefficients. Viral families with an average relative abundance below 0.01% were excluded. The transformed matrix was used to examine viral family dominance across phyla through a heatmap (Figure 4B). PCA was also performed, but the biplot representation of the results was not visually intuitive because of the large amount of data points and different groups defined by phylum and habitat in this dataset. The first two principal components accounted for 23% of the total variance, indicating a limited correlation among the *clr* coefficients of the 30 viral families studied.

Hierarchical clustering analysis revealed the presence of three clusters of viral families that were distinguishable with considerable resolution (Figure 4B). The first cluster consisted of 13 superabundant, ubiquitous viral families (*Baculo*-, *Flavi*-, *Hepe*-, *Irido*-, *Marseille*-, *Mimi*-, *Mito*-, *Partiti*-, *Phycodna*-, *Pitho*-, *Polydnaviriformi*-, *Pox-*, and *Retroviridae*) showing positive *clr* coefficients for all phyla (Figure 4B). Within this cluster, eight of the families harbored viruses with dsDNA genomes (*Baculo*-, *Irido*-, *Marseille*-, *Mimi*-, *Phycodna*-, *Pitho*-, *Polydnaviriformi*-, and *Poxviridae*), with a notable presence of NCLDV families. In the analysis, all NCLDV families except *Ascoviridae* clustered together in this group. Furthermore, this cluster was the one with the highest distance to the other two clusters.

A second cluster contained less abundant families with values of *clr* coefficients oscillating around zero (*Alloherpes*-, *Asco*-, *Nairo*-, *Herpes*-, *Hytrosa-*, *Nudi*-, and *Picornaviridae*), signifying their dominance within the virome only for specific phyla (Figure 4B). DNA virus families are again more frequent here (five out of seven) and the acknowledged host ranges of these families include invertebrates in all cases except for *Herpes-* and *Alloherpesviridae*.

The last cluster is constituted by even rarer viral families that were only found to be dominant in the viromes of one or very few phyla: *Adeno*-, *Adinto*-, *Adoma*-, *Betaflexi*-, *Dicistro*-, *Iflavi*-, *Marna*-, *Nima*-, *Polycipi*-, and *Potyviridae* (Figure 4B). Among the ten viral families included in this cluster, six of them were families of RNA viruses. The two only families of exclusively plant and/or fungi-infecting viruses of this 30 families dataset were found in this cluster: *Betaflexi-* and *Potyviridae*. In both cases, sequences similar to these families were identified predominantly in phyla of marine invertebrates, where the occurrence of these viruses has not been confirmed. In the case of *Betaflexiviridae*, most hits pointed to the occurrence of viral replicases and RNA-dependent RNA polymerases within the samples. Meanwhile, most of the hits of *Potyviridae* were related to an inosine triphosphate pyrophosphatase (ITPase) protein, which is only present in two potyvirus species (Tomlinson et al., 2019). This protein shows significant sequence homology to ITPase proteins of plants, and, to a lesser extent, in invertebrates. However, hits to coat proteins of potyviruses were also reported in several samples. True viral presence of both families among the samples seems likely, although the abundance of potyviruses was probably overestimated. Interestingly, *Betaflexiviridae*-like sequences were also identified in different transcriptomes of marine mollusk samples by a recent study (Rey-Campos et al., 2023).

Unlike the clustering observed for viral families, the separation of phyla into clusters was less obvious. Interpreting the clusters proved more challenging, not only because of the clustering’s low resolution, but also for the apparent lack of consistent patterns within the clusters based on the phylogenetic or habitat affinities between the phyla. One of the groups with a greater distance to the rest was the one formed by Arthropoda and Onychophora, two phyla that are closely related phylogenetically (Dunn et al., 2008). This suggests that their viromes have similar compositions at the family level. For instance, these two were the only phyla where sequences of *Iflaviridae*, a family of insect-infecting RNA viruses, were found with a high relative abundance.

PERMANOVA models were fitted in order to investigate the potential impact that the phylum and the habitat of an invertebrate sample could have on the composition of its virome. The dataset was restricted to phyla and habitats with 15 or more samples for the analysis. In both one-way models, it was seen that virome compositions were significantly influenced by the host’s phylum and habitat (*P* < 0.05). Nonetheless, the effect of the host’s phylum on the compositions was nearly six times greater than that of habitat (see Table S6 at [Supplementary_files]). It has to be taken into account that the habitat effect could have been underestimated in this study because of the lack of a more comprehensive set of habitat categories. The phylum factor alone accounted for approximately 26% of the variance in the Euclidean distances matrix.

In the additive two-way model, regression parameters associated to both factors remained significant, suggesting that collinearity, although present, does not totally neglect the separate effect of both factors in shaping the viromes. Furthermore, the model considering phylum and habitat interaction showed that even the interaction had significant regression parameters. Nevertheless, the slight improvement in goodness of fit did not offset the increase in model complexity of the two-way models, since the model with phylum as its single covariate was the one that minimized AIC. It was checked that similar results overall were obtained when using the Bray-Curtis dissimilarities of the raw relative abundances matrix, which is a more traditional approach.

However, it’s important to note that the rejection of the null hypothesis in PERMANOVA could also arise from varying within-group dispersions in the case of unbalanced designs (Anderson & Walsh, 2013). The result of PERMDISP showed that within-group dispersions were indeed heterogeneous between phyla (*P* < 0.05). Virome composition variation was more thoroughly examined using a procedure based on the analysis of the distances to the centroid, *i.e.*, the “average virome”, in each phylum. The strategy of fitting an ANOVA model and performing a Tukey’s *post hoc* test was also used for this variable and showed that distances to centroid were significantly different between some phyla (*P* < 0.05). Interestingly, most of the phyla exhibiting extreme distances to centroid have already been mentioned for having an exceptional behavior in terms of ⍺-diversity. Bryozoans appeared to have the most consistently similar virome compositions, closely followed by ctenophores. Conversely, mollusks showed the greatest mean distance to the centroid, together with xenacoelomorphs and sponges (Figure 3D).

The habitat variability between phyla does not appear to be the primary driver of these observed differences, as phyla with both diverse and singular habitats were represented at both extremes. One potential influencing factor for these findings could be the phylogenetic proximity among species within the same phylum. This study treats species as independent observations of their respective phylum, disregarding their membership in the same genus, family, or order. This is a limitation of this study, since the phylogenetic constraints underlying species relatedness within phyla were not taken into consideration.

Additionally, the viromes of Arthopoda, Onychophora and Xenacoelomorpha exhibit notable variability in their distances to the phylum centroid. This indicates that within these phyla, certain samples are positioned very close to the centroid, while others are located considerably farther away (Figure 3D). That is to say, these phyla tend to show a high variability in their virome compositions, not only as the different viral families that can be found in these viromes, but also in what relative abundances they are found.

A pairwise algorithm with bootstrap resampling and simultaneous PERMANOVA and PERMDISP testing was implemented to further investigate if virome compositions were different between phyla. To ensure balanced group sizes, the sample count per phylum was standardized to 15. The results highlighted that a significant majority of phyla pairs exhibited dissimilar centroid locations in most iterations (see Figure S2B at [Supplementary_files]), even if a *P*-value adjustment method was employed to correct for multiple pairwise comparisons. Moreover, equivalence of centroid locations was observed as the most frequent result through the iterations only for a limited number of phyla pairs. Surprisingly, heterogeneous within-group dispersions were not found to be the most frequent result in any case, in contrast to earlier findings using the whole dataset.

The results of this matrix were condensed into a network to allow a visual interpretation of the results (Figure 4C). Curiously, all phyla remained connected to at least some other phylum, meaning that their virome composition centroids were not considered significantly different to each other. This resulted in a continuous network, where no phylum had an outlying virome composition that differed from the rest. However, there were some cases of phyla with few and weak connections, such as Ctenophora, Echinodermata and Platyhelminthes. In contrast, Mollusca was the phylum with the highest number of connections, some of which were strong, meaning that the “average virome” of mollusks was not distinguishable from that of some other phyla.

Some of the strongest similarities between virome compositions that can be observed in the network, can also be identified in the heatmap (Figure 4B), namely Arthropoda- Onychophora, Mollusca-Annelida, Kinorhyncha-Porifera, and Rotifera-Nematoda. Viromes of Arthropoda only appear to be closely similar to those of Onychophora, which also shares a rather pronounced centroid homology with Annelida, all three being phyla dominated by terrestrial species. In the case of Arthropoda and Onychophora, both belong to the clade of Panarthropoda, alongside Tardigrada (Wu et al., 2023 and Figure 1C), which could explain the remarkable degree of similarity observed between their virome compositions. Unfortunately, Tardigrada could not be included in this part of the analysis to test this hypothesis further because of its low sample count. Likewise, the reported similarity between the viromes of Mollusca and Annelida could be attributed to their phylogenetic relatedness, since both group within a subclade under the superphylum Lophotrochozoa (Figure 1C) (Kocot et al., 2017). Moreover, the average viromes of Nemertea showed weak resemblance to those of the rest phyla in this subclade, while the similarity between the viromes of Mollusca and Bryozoa, another phylum of this subclade, was one of the highest (Figure 4C). The hypothesis behind this phenomenon of closely related phyla having equivalent viromes is that viromes may have coevolved with their hosts. Viromes of these phyla are probably less influenced by changes in the host’s habitat or lifestyle during evolution. Nonetheless, this explanation seems applicable only for the virome composition similarities of a limited number of phyla.

In other cases, such as the reported similarity of virome compositions between Kinorhyncha and Porifera or Nemertea and Xenacoelomorpha, the explanation could not be found in the phylogenetic relationships between the phyla. Rather, these phyla appeared to have similar habitats among their samples. However, it is unknown why the viromes of these specific phyla were found to be so similar, given that many other phyla have apparently similar habitats. For instance, it has been described that kinorhynchs can be found inhabiting sponges, although this is not thought to be the norm and they are generally interstitial organisms (Higgins, 1978; Neuhaus & Higgins, 2002). Therefore, a more comprehensive analysis of lifestyles, diet, and associations with other organisms would be needed in this regard.

The reason behind some other phyla having notable similarities across their average virome compositions, such as Kinorhyncha and Mollusca, Bryozoa and Kinorhyncha or Nematoda and Rotifera, remains uncertain, since these specific pairs of phyla do not exhibit any close phylogenetic or ecological affinity. But in summary, our findings align with the results of Shi et al. (2016). They underscore that the assumption of virus-host co-divergence cannot be inferred from all cases where viromes exhibit similarities across different phyla, since transmission of viruses between divergent host taxa has been frequent throughout the evolutionary history of viruses (Dolja & Koonin, 2018).

## Conclusions

A read taxonomic classification pipeline for RNA and DNA viruses was applied to more than 400 RNA-Seq datasets of invertebrates comprising 31 phyla. Overall, the estimated viromes were not widely variable in terms of ⍺-diversity, although significant differences in the means could be found between the phyla. Additionally, viromes were consistently dominated by a set of ubiquitous families, the abundance of which was likely overestimated due to the misclassification of host sequences as viral, as such is the case for families like *Retroviridae*, *Flaviviridae* or *Herpesviridae*. The structure of the invertebrate-virus bipartite network showed that its interactions were highly nested, which hints to a shared core virome across the Animal Tree of Life. In the majority of the most abundant viral families found, their acknowledged host ranges in the literature included invertebrates. However, the confirmed host ranges of these families are limited to overrepresented phyla such as Arthropoda or Mollusca and the presence of most of these families has still to be confirmed for the most understudied phyla. Consequently, virus sequences in the reference databases are also biased toward these relevant phyla.

Further improvements in the resolution of the Animal Tree of Life could help improve the interpretation of invertebrate viromes, since many nodes remain unresolved and changes in the phylogeny of animal phyla keep occurring to this date. However, evidence shows that the host’s phylum has a predominant role in shaping its virome composition, rather than its habitat. A main limitation for this observation could be the effect that differential expression of host genes that had been erroneously classified as viral could have had between the phyla. Also, it is important to note that the reconstruction of the host’s habitat was based on the taxonomy of the host rather than the sampling source due to the lack of metadata in many cases. Consequently, the host’s habitat affiliation was limited to a very generic set of habitat categories.

As a concluding remark, comparability to other studies has remained limited due to the lack of studies analyzing together the RNA and DNA viromes of invertebrates at the phylum level. Additionally, ecological community structure of invertebrate-virus interactions at the phylum and family levels, respectively, and the systematic review of similarities between the virome compositions of the different phyla had not yet been published. Finally, we recommend that similar studies implement a more intensive filtering of host sequences that is not limited to rRNA transcripts, although a trade-off will always exist between false positives and false negatives when classifying host sequences as viral and excluding transcripts encoded by the viral genome with true homology to eukaryotic genes, respectively.

## Acknowledgements

PA was supported by fellowship JAEIntro-2021-LifeHub-10 from CSIC. AB was supported by a post-doctoral fellowship from Foundation pour la Recherche Mèdicale (grant number SPF202110014092). RF acknowledges support from the following sources of funding: Ramón y Cajal fellowship (grant agreement no. RYC2017-22492 funded by MCIN/AEI /10.13039/501100011033 and ESF ‘Investing in your future’), the Agencia Estatal de Investigación (project PID2019-108824GA-I00 funded by MCIN/AEI/10.13039/501100011033), the European Research Council (this project has received funding from the European Research Council (ERC) under the European’s Union’s Horizon 2020 research and innovation programme (grant agreement no. 948281)), the Human Frontier Science Program (grant no. RGY0056/2022) and the Secretaria d’Universitats i Recerca del Departament d’Economia i Coneixement de la Generalitat de Catalunya (AGAUR 2021-SGR00420). AR acknowledges support from: Ramón y Cajal fellowship (grant agreement no. RYC2018-024247-I funded by MCIN/AEI /10.13039/501100011033 and ESF ‘Investing in your future’), the Agencia Estatal de Investigación (project PID2019-105769GB- I00 funded by MCIN/AEI/10.13039/501100011033), an internal grant from CSIC (PIE- 202030E006), and a 2021-2022 BiodivProtect joint call for research proposals, under the Biodiversa+ Partnership co-funded by the European Commission, and with the funding organisations ‘Fundación Biodiversidad’ and FORMAS. RF, AR and SFE also acknowledge support and funding from LifeHUB CSIC (PIE-202120E047).

## Supplementary materials

### Pre-processing of raw reads results

Deduplication produced a very uneven filtering of reads between samples, ranging from almost no duplicates found to a decline of approximately 90% of reads in the most extreme cases. On average, approximately 32% of reads were filtered out for being exact duplicates of others. Contrary to what would be expected, the proportion of duplicates found in each sample showed no relation to its library size. So, sequencing depth did not produce a significantly higher proportion of exact duplicates.

Removal of rRNA and *COX* reads from the subset of non-duplicated reads yielded a further decline in the number of reads of 56% on average. In this case, there was less variability and the large majority of samples lost 50 to 60% of the already non-duplicated reads. Only in five samples a reduction of more than 80% of the reads was noted and in no case larger than 90%. This step involved the most intense removal of reads among the pre-processing methods. However, an even larger proportion (> 90%) of rRNA reads could be expected in invertebrate transcriptomes (Kumar et al., 2012). Prior deduplication could have had an effect in preferentially excluding reads of superabundant RNA molecules such as rRNA. Furthermore, prokaryotic rRNA was not targeted in this study.

Last, the step of quality filtering yielded a reduction of less than 10% of the already non-duplicated, non-rRNA reads in more than 75% of the samples. In an outlying case, quality filtering led to a loss of 90% of the remaining reads in the sample of *Rhynchomesostoma sp.* (Platyhelminthes).

In all cases, at least 45% of the total raw reads were discarded following the application of the three pre-processing methods. On average, only 27% of the original reads were preserved and the interquartile range lied between 20% and 36%, which represents the central 50% of the distribution of preserved reads. With the exception of a single sample, the remaining number of reads for all samples was at least one million while half of the samples had more than 15 million reads left after pre-processing (Figure 1A).

**Figure S1.**
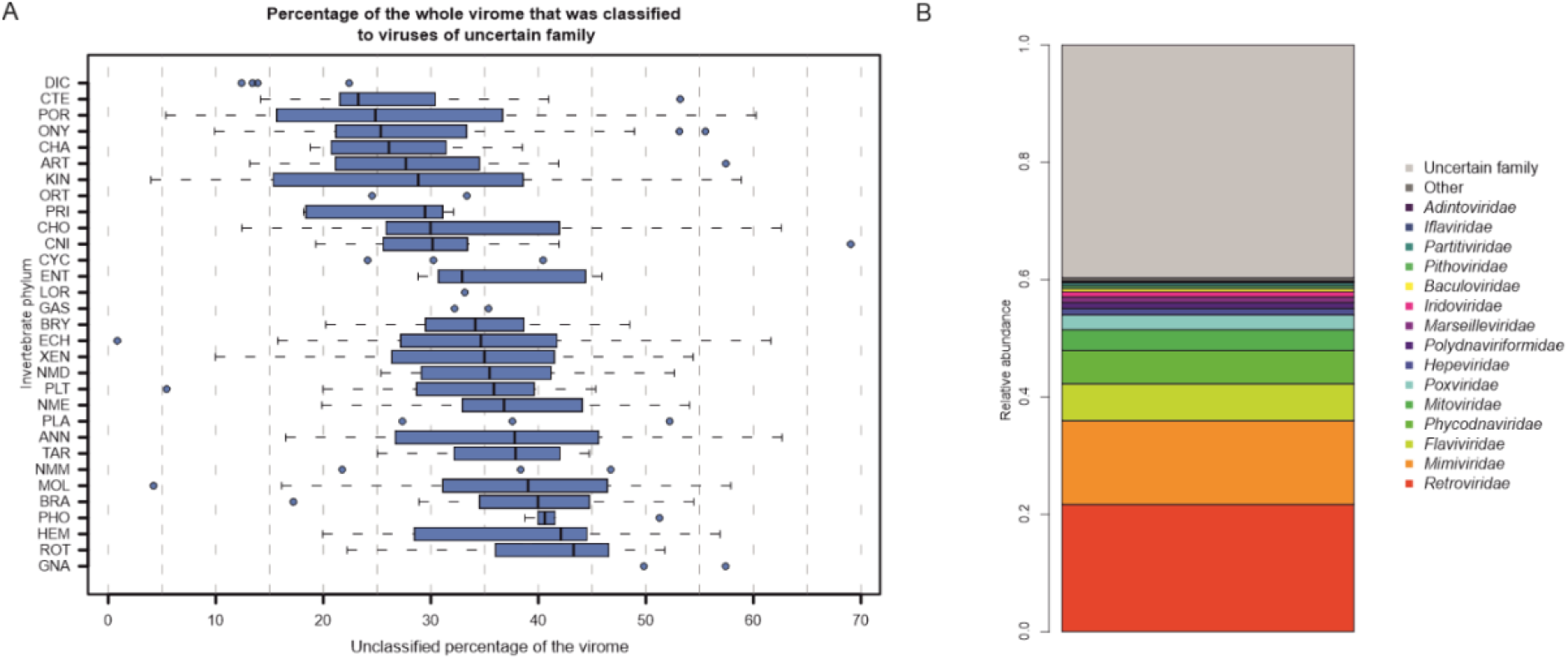
**A.** Distribution by phylum of the percentage of the virome that was classified to viruses of uncertain family. Boxes are not drawn for phyla with less than 5 samples. Full phylum names can be found in Table S2 at [Supplementary_files]. **B.** Compositional barplot of the global virome. Phage families were excluded. All samples were given the same weight when aggregating the relative abundances vectors of all 417 samples.

**Figure S2.**
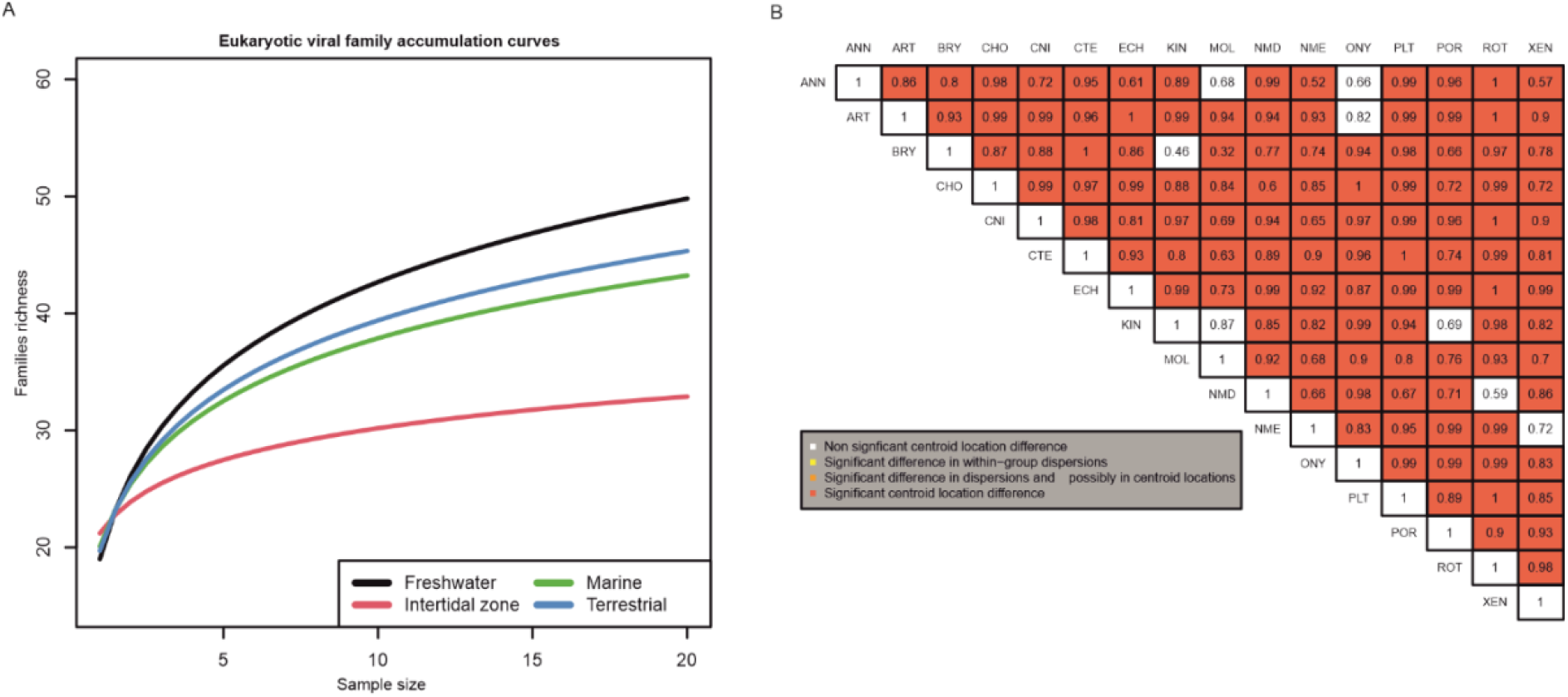
**A.** Eukaryotic viral families accumulation curves by habitat. 100 resampling iterations were run at each sample size to avoid bias. Brackish habitat was excluded. **B.** Hemimatrix showing the results of the pairwise PERMANOVA analysis with bootstrap. Cell color represents the most frequent combination of test results for each pair of phyla (red: H_1_ of PERMANOVA + H_0_ of PERMDISP; orange: H_1_ of PERMANOVA + H_1_ of PERMDISP; yellow: H_0_ of PERMANOVA + H_1_ of PERMDISP; white: H_0_ of PERMANOVA + H_0_ of PERMDISP. The values represent the proportions of iterations where the most frequent result is found.

**Table S1**. Number of species with one or more eligible accessions (see criteria in materials and methods) in the SRA archive by phylum. (CSV)

**Table S2**. Meaning of the phylum abbreviations used in Figures 1B, 2C, 3A-D, 4A-C, S1A and S2B. (CSV)

**Table S3.** Count and percentage of samples where the presence of at least one read of each viral family of eukaryotes was recorded.

**Table S4**. Overview of statistics and other results obtained for top 30 viral families by mean relative abundance. (CSV)

**Table S5**. Summary of statistics for the comparison between observed community structure metrics and their simulated under null models counterpart. (CSV)

**Table S6**. Comparison of PERMANOVA model statistics between formulae using different combinations of covariates. (CSV)

## Data and scripts availability

Supplementary Files can be downloaded from https://git.csic.es/sfelenalab/invertebrates-virome/-/tree/main/Supplementary%20files. All scripts used for the selection of samples, the read taxonomic classification pipeline and the statistical analysis, as well as the generation of figures presented in this report, can be found at https://git.csic.es/sfelenalab/invertebrates-virome under the directory “Data and Scripts”. Intermediate datasets, in addition to the main data table in CSV format, can be found alongside the scripts. The code for the Shiny app is also available at the same Git repository under the directory “Shiny app”, allowing the app to be run locally.

## References

Ahyong S, et al. 2023. World Register of Marine Species. Available from https://www.marinespecies.org at VLIZ. Accessed 2023-09-19. 10.14284/170

Almeida-Neto M, Guimarães P, Guimarães Jr PR, Loyola RD. and Ulrich W. 2008. A consistent metric for nestedness analysis in ecological systems: reconciling concept and measurement. Oikos 117(8):1227–1239. 10.1111/j.0030-1299.2008.16644.x

Anderson MJ. 2006. Distance-based tests for homogeneity of multivariate dispersions. Biometrics 62(1): 245–253. 10.1111/j.1541-0420.2005.00440.x

Anderson MJ. 2001. A new method for non-parametric multivariate analysis of variance. Austral Ecology 26(1): 32–46. 10.1111/J.1442-9993.2001.01070.PP.X

Anderson MJ, and Walsh DCI. 2013. PERMANOVA, ANOSIM, and the Mantel test in the face of heterogeneous dispersions: What null hypothesis are you testing?. Ecological Monographs 83(4): 557–574. 10.1890/12-2010.1

Blüthgen N, Menzel F, and Blüthgen N. 2006. Measuring specialization in species interaction networks. BMC ecology 6,9. 10.1186/1472-6785-6-9

Bolger AM, Lohse M, and Usadel B. 2014. Trimmomatic: a flexible trimmer for Illumina sequence data. Bioinformatics 30(15): 2114–2120. 10.1093/bioinformatics/btu170

Boyer M, Yutin N, Pagnier I, Barrassi L, Fournous G, Espinosa L, Robert C, Azza S, Sun S, Rossmann MG, Suzan-Monti M, La Scola B, Koonin EV, and Raoult D. 2009. Giant Marseillevirus highlights the role of amoebae as a melting pot in emergence of chimeric microorganisms. Proceedings of the National Academy of Sciences of the United States of America 106(51): 21848–21853. 10.1073/pnas.0911354106

Breitbart M, and Rohwer F. 2005. Here a virus, there a virus, everywhere the same virus?. Trends in microbiology 13(6): 278–284. 10.1016/j.tim.2005.04.003

Brüwer JD, and Voolstra CR. 2018. First insight into the viral community of the cnidarian model metaorganism Aiptasia using RNA-Seq data. PeerJ 6: e4449. 10.7717/peerj.4449

Canuti M, Large G, Verhoeven JTP, and Dufour SC. 2022. A Novel Iridovirus Discovered in Deep-Sea Carnivorous Sponges. Viruses 14(8): 1595. 10.3390/v14081595

Cobbin JC, Charon J, Harvey E, Holmes EC, and Mahar JE. 2021. Current challenges to virus discovery by meta-transcriptomics. Current opinion in virology 51: 48–55. 10.1016/j.coviro.2021.09.007

Cross ST, Maertens BL, Dunham TJ, Rodgers CP, Brehm AL, Miller MR, Williams AM, Foy BD, and Stenglein MD. 2020. Partitiviruses Infecting Drosophila melanogaster and Aedes aegypti Exhibit Efficient Biparental Vertical Transmission. Journal of virology 94(20): e01070–20. 10.1128/JVI.01070-20

Csardi G, and Nepusz T. 2006. The igraph software package for complex network research. InterJournal, Complex Systems 1695. https://igraph.org.

de Souza, FG, Abrahão JS, and Rodrigues RAL. 2021. Comparative Analysis of Transcriptional Regulation Patterns: Understanding the Gene Expression Profile in *Nucleocytoviricota*. Pathogens 10(8): 935. 10.3390/pathogens10080935

Dolja VV, and Koonin EV. 2018. Metagenomics reshapes the concepts of RNA virus evolution by revealing extensive horizontal virus transfer. Virus research 244: 36–52. 10.1016/j.virusres.2017.10.020

Dormann C, Frund J, Blüthgen N, and Gruber B. 2009. Indices, Graphs and Null Models: Analyzing Bipartite Ecological Networks. Open Journal of Ecology 2: 7–24. 10.2174/1874213000902010007

Dufour F, Bertrand L, Pearson A, Grandvaux N, and Langelier Y. 2011. The ribonucleotide reductase R1 subunits of herpes simplex virus 1 and 2 protect cells against poly(I · C)- induced apoptosis. Journal of virology 85(17): 8689–8701. 10.1128/JVI.00362-11

Dunn CW, Hejnol A, Matus DQ, Pang K, Browne WE, Smith SA, Seaver E, Rouse GW, Obst M, Edgecombe GD, Sørensen MV, Haddock SH, Schmidt-Rhaesa A, Okusu A, Kristensen RM, Wheeler WC, Martindale MQ, and Giribet G. 2008. Broad phylogenomic sampling improves resolution of the animal tree of life. Nature 452(7188): 745–749. 10.1038/nature06614

Farzad R, Ha AD, and Aylward FO. 2022. Diversity and genomics of giant viruses in the North Pacific Subtropical Gyre. Frontiers in microbiology 13: 1021923. 10.3389/fmicb.2022.1021923

Feng Y, Krueger EN, Liu S, Dorman K, Bonning BC, and Miller WA. 2017. Discovery of Known and Novel Viral Genomes in Soybean Aphid by Deep Sequencing. Phytobiomes Journal 1: 36–45. 10.1094/PBIOMES-11-16-0013-R

Fox J, and Weisberg S. 2019. An R Companion to Applied Regression (Third edition). Thousand Oaks, CA: Sage. Retrieved from https://socialsciences.mcmaster.ca/jfox/Books/Companion/

García-Bonilla E, Chaves-Moreno D, Riaño-Pachón D, Terán W, Acosta A, and Junca H. 2021. The Hologenome of Haliclona fulva (Porifera, Demospongiae) Reveals an Abundant and Diverse Viral Community. Frontiers in Marine Science 8: 1632. 10.3389/fmars.2021.736817

Gáspár G, De Clercq E, and Neyts J. 2002. Human herpesvirus 8 gene encodes a functional thymidylate synthase. Journal of virology 76(20): 10530–10532. 10.1128/jvi.76.20.10530-10532.2002

Gilbert C, and Belliardo C. 2022. The diversity of endogenous viral elements in insects. Current opinion in insect science 49: 48–55. 10.1016/j.cois.2021.11.007

Gloor GB, Macklaim JM, Pawlowsky-Glahn V, and Egozcue JJ. 2017. Microbiome Datasets Are Compositional: And This Is Not Optional. Frontiers in microbiology 8: 2224. 10.3389/fmicb.2017.02224

Grasis JA, Lachnit T, Anton-Erxleben F, Lim YW, Schmieder R, Fraune S, Franzenburg S, Insua S, Machado G, Haynes M, Little M, Kimble R, Rosenstiel P, Rohwer FL, and Bosch TC. 2014. Species-specific viromes in the ancestral holobiont Hydra. PloS one 9(10): e109952. 10.1371/journal.pone.0109952

Graves S, Piepho H-P, Selzer L, and Dorai-Raj S. 2023. multcompView: Visualizations of Paired Comparisons (Version 0.1-9) [Software manual]. https://CRAN.R-project.org/package=multcompView

Gudenkauf BM, and Hewson I. 2016. Comparative metagenomics of viral assemblages inhabiting four phyla of marine invertebrates. Frontiers in Marine Science 3: 23. 10.3389/fmars.2016.00023

Ha AD, Moniruzzaman M, and Aylward FO. 2021. High Transcriptional Activity and Diverse Functional Repertoires of Hundreds of Giant Viruses in a Coastal Marine System. mSystems 6(4): e0029321. 10.1128/mSystems.00293-21

Harvey E, and Holmes EC. 2022. Diversity and evolution of the animal virome. Nat Rev Microbiol 20(6): 321–334. 10.1038/s41579-021-00665-x

Herranz M, Stiller J, Worsaae K, and Sørensen MV. 2022. Phylogenomic analyses of mud dragons (Kinorhyncha). Molecular phylogenetics and evolution 168: 107375. 10.1016/j.ympev.2021.107375

Hewson I, Aquino CA, and DeRito CM. 2020. Virome Variation during Sea Star Wasting Disease Progression in *Pisaster ochraceus* (Asteroidea, Echinodermata). Viruses 12(11): 1332. 10.3390/v12111332

Holt CC, Boscaro V, Van Steenkiste NWL, Herranz M, Mathur V, Irwin NAT, Buckholtz G, Leander BS, and Keeling PJ. 2022. Microscopic marine invertebrates are reservoirs for cryptic and diverse protists and fungi. Microbiome 10(1): 161. 10.1186/s40168-022-01363-3

Huang HJ, Ye ZX, Wang X, Yan XT, Zhang Y, He YJ, Qi YH, Zhang XD, Zhuo JC, Lu G, Lu JB, Mao QZ, Sun ZT, Yan F, Chen JP, Zhang CX, and Li JM. 2021. Diversity and infectivity of the RNA virome among different cryptic species of an agriculturally important insect vector: whitefly Bemisia tabaci. NPJ biofilms and microbiomes 7(1): 43. 10.1038/s41522-021-00216-5

Jacquat AG, Ulla SB, Debat HJ, Muñoz-Adalia EJ, Theumer MG, Pedrajas MDG, and Dambolena JS. 2022. An in silico analysis revealed a novel evolutionary lineage of putative mitoviruses. Environmental microbiology 24(12): 6463–6475. 10.1111/1462-2920.16202

Juravel K, Porras L, Höhna S, Pisani D, and Wörheide G. 2023. Exploring genome gene content and morphological analysis to test recalcitrant nodes in the animal phylogeny. PloS one 18(3): e0282444. 10.1371/journal.pone.0282444

Kocot KM, Struck TH, Merkel J, Waits DS, Todt C, Brannock PM, Weese DA, Cannon JT, Moroz LL, Lieb B, and Halanych KM. 2017. Phylogenomics of Lophotrochozoa with Consideration of Systematic Error. Systematic biology 66(2): 256–282. 10.1093/sysbio/syw079

Kondo H, Fujita M, Hisano H, Hyodo K, Andika IB, and Suzuki N. 2020. Virome Analysis of Aphid Populations That Infest the Barley Field: The Discovery of Two Novel Groups of Nege/Kita-Like Viruses and Other Novel RNA Viruses. Frontiers in microbiology 11: 509. 10.3389/fmicb.2020.00509

Koonin, E. V., & Yutin, N. (2010). Origin and evolution of eukaryotic large nucleo- cytoplasmic DNA viruses. Intervirology, 53(5), 284–292. 10.1159/000312913

Koonin EV, Senkevich TG, and Dolja VV. 2006. The ancient Virus World and evolution of cells. Biology direct 1: 29. 10.1186/1745-6150-1-29

Kopylova E, Noé L, and Touzet H. 2012. SortMeRNA: fast and accurate filtering of ribosomal RNAs in metatranscriptomic data. Bioinformatics 28(24): 3211–3217. 10.1093/bioinformatics/bts611

Kumar N, Creasy T, Sun Y, Flowers M, Tallon LJ, and Dunning Hotopp JC. 2012. Efficient subtraction of insect rRNA prior to transcriptome analysis of Wolbachia-Drosophila lateral gene transfer. BMC research notes 5: 230. 10.1186/1756-0500-5-230

Laumer CE, Fernández R, Lemer S, Combosch D, Kocot KM, Riesgo A, Andrade SCS, Sterrer, W, Sørensen MV, and Giribet G. 2019. Revisiting metazoan phylogeny with genomic sampling of all phyla. *Proceedings*, Biological sciences 286(1906): 20190831. 10.1098/rspb.2019.0831

Laumer CE, Gruber-Vodicka H, Hadfield MG, Pearse VB, Riesgo A, Marioni JC, and Giribet G. 2018. Support for a clade of Placozoa and Cnidaria in genes with minimal compositional bias. eLife 7: e36278. 10.7554/eLife.36278

Lewandowska M, Hazan Y, and Moran Y. 2020. Initial Virome Characterization of the Common Cnidarian Lab Model *Nematostella vectensis*. Viruses 12(2): 218. 10.3390/v12020218

Li Y, Shen XX, Evans B, Dunn CW, and Rokas A. 2021. Rooting the Animal Tree of Life. Molecular biology and evolution 38(10): 4322–4333. 10.1093/molbev/msab170

Lu TM, Kanda M, Satoh N, and Furuya H. 2017. The phylogenetic position of dicyemid mesozoans offers insights into spiralian evolution. Zoological letters 3: 6. 10.1186/s40851-017-0068-5

Marlétaz F, Peijnenburg KTCA, Goto T, Satoh N, and Rokhsar DS. 2019. A New Spiralian Phylogeny Places the Enigmatic Arrow Worms among Gnathiferans. Current biology 29(2): 312–318.e3. 10.1016/j.cub.2018.11.042

Mayer D, Thayer TM, Hofmann MA, and Tratschin JD. 2003. Establishment and characterisation of two cDNA-derived strains of classical swine fever virus, one highly virulent and one avirulent. Virus research 98(2): 105–116. 10.1016/j.virusres.2003.08.020

Meng F, Ding M, Tan Z, Zhao Z, Xu L, Wu J, He B, and Tu C. 2019. Virome analysis of tick- borne viruses in Heilongjiang Province, China. Ticks and tick-borne diseases 10(2): 412–420. 10.1016/j.ttbdis.2018.12.002

Menzel P, Ng KL, and Krogh A. 2016. Fast and sensitive taxonomic classification for metagenomics with Kaiju. Nature Communications 7: 11257. 10.1038/ncomms11257

Moureau G, Cook S, Lemey P, Nougairede A, Forrester NL, Khasnatinov M, Charrel RN, Firth AE, Gould EA, and de Lamballerie X. 2015. New insights into flavivirus evolution, taxonomy and biogeographic history, extended by analysis of canonical and alternative coding sequences. PloS one 10(2): e0117849. 10.1371/journal.pone.0117849

Mouritsen KN, and Poulin R. 2002. Parasitism, community structure and biodiversity in intertidal ecosystems. Parasitology 124(7): 101–117. 10.1017/s0031182002001476

Neri U, Wolf YI, Roux S, Camargo AP, Lee B, Kazlauskas D, Chen IM, Ivanova N, Zeigler Allen L, Paez-Espino D, Bryant DA, Bhaya D, RNA Virus Discovery Consortium, Krupovic M, Dolja VV, Kyrpides NC, Koonin EV, and Gophna U. 2022. Expansion of the global RNA virome reveals diverse clades of bacteriophages. Cell 185(21): 4023–4037.e18. 10.1016/j.cell.2022.08.023

Neuhaus B, and Higgins RP. 2002. Ultrastructure, biology, and phylogenetic relationships of kinorhyncha. Integrative and comparative biology 42(3): 619–632. 10.1093/icb/42.3.619

Newman MEJ. 2006. Modularity and community structure in networks. Proceedings of the National Academy of Sciences 103(23): 8577–8582. 10.1073/pnas.0601602103

Oksanen J, et al. 2022. vegan: Community Ecology Package (Version 2.6-4) [Software manual]. https://CRAN.R-project.org/package=vegan

Olendraite I, Brown K, and Firth AE. 2023. Identification of RNA Virus-Derived RdRp Sequences in Publicly Available Transcriptomic Data Sets. Molecular biology and evolution 40(4): msad060. 10.1093/molbev/msad060

Palarea-Albaladejo J, and Martín-Fernández JA. 2015. zCompositions — R package for multivariate imputation of left-censored data under a compositional approach. Chemometrics and Intelligent Laboratory Systems 143: 85–96. 10.1016/j.chemolab.2015.02.019

Parry R, and Asgari S. 2019. Discovery of Novel Crustacean and Cephalopod Flaviviruses: Insights into the Evolution and Circulation of Flaviviruses between Marine Invertebrate and Vertebrate Hosts. Journal of virology 93(14): e00432–19. 10.1128/JVI.00432-19

Pascelli C, Laffy PW, Botté E, Kupresanin M, Rattei T, Lurgi M, Ravasi T, and Webster N. S. 2020. Viral ecogenomics across the Porifera. Microbiome 8(1): 144. 10.1186/s40168-020-00919-5

Perez-Riverol Y, Zorin A, Dass G, Vu MT, Xu P, Glont M, Vizcaíno JA, Jarnuczak AF, Petryszak R, Ping P, and Hermjakob H. 2019. Quantifying the impact of public omics data. Nature communications 10(1): 3512. 10.1038/s41467-019-11461-w

Porter AF, Shi M, Eden JS, Zhang YZ, and Holmes EC. 2019. Diversity and Evolution of Novel Invertebrate DNA Viruses Revealed by Meta-Transcriptomics. Viruses 11(12): 1092. 10.3390/v11121092

Raoult D, Audic S, Robert C, Abergel C, Renesto P, Ogata H, La Scola B, Suzan M, and Claverie JM. 2004. The 1.2-megabase genome sequence of Mimivirus. Science 306(5700): 1344–1350. 10.1126/science.1101485

Rey-Campos M, González-Vázquez LD, Novoa B, and Figueras A. 2023. Metatranscriptomics unmasks Mollusca virome with a remarkable presence of rhabdovirus in cephalopods. Frontiers in Marine Science 10: 1209103. 10.3389/fmars.2023.1209103

Robert P Higgins. 1978. *Echinoderes gerardi n. sp.* and *E. riedli* (Kinorhyncha) from the Gulf of Tunis. Transactions of the American Microscopical Society 97(2): 171–180. 10.2307/3225589

Rosani U, Gaia M, Delmont TO, and Krupovic M. 2023. Tracing the invertebrate herpesviruses in the global sequence datasets. Frontiers in Marine Science 10: 1159754. 10.3389/fmars.2023.1159754

Schultz DT, Haddock SHD, Bredeson JV, Green RE, Simakov O, and Rokhsar DS. 2023. Ancient gene linkages support ctenophores as sister to other animals. Nature 618: 110–117. 10.1038/s41586-023-05936-6

Shi C, Beller L, Deboutte W, Yinda KC, Delang L, Vega-Rúa A, Failloux AB, and Matthijnssens J. 2019. Stable distinct core eukaryotic viromes in different mosquito species from Guadeloupe, using single mosquito viral metagenomics. Microbiome 7(1),:121. 10.1186/s40168-019-0734-2

Shi C, Zhao L, Atoni E, Zeng W, Hu X, Matthijnssens J, Yuan Z, and Xia H. 2020. Stability of the Virome in Lab- and Field-Collected Aedes albopictus Mosquitoes across Different Developmental Stages and Possible Core Viruses in the Publicly Available Virome Data of Aedes Mosquitoes. mSystems 5(5): e00640–20. 10.1128/mSystems.00640-20

Shi M, Lin XD, Tian JH, Chen LJ, Chen X, Li CX, Qin XC, Li J, Cao JP, Eden JS, Buchmann J, Wang W, Xu J, Holmes EC, and Zhang YZ. 2016. Redefining the invertebrate RNA virosphere. Nature 540(7634): 539–543. 10.1038/nature20167

Simmonds P, Adams MJ, Benkő M, Breitbart M, Brister JR, Carstens EB, Davison AJ, Delwart E, Gorbalenya AE, Harrach B, Hull R, King AM, Koonin EV, Krupovic M, Kuhn JH, Lefkowitz EJ, Nibert ML, Orton R, Roossinck MJ, Sabanadzovic S, Sullivan MB, Suttle CA, Tesh RB, van der Vlugt RA, Varsani A, and Zerbini FM. 2017a. Consensus statement: Virus taxonomy in the age of metagenomics. Nature reviews Microbiology 15(3): 161–168. 10.1038/nrmicro.2016.177

Simmonds P, et al. 2023. Four principles to establish a universal virus taxonomy. PLoS Biology 21(2): e3001922. 10.1371/journal.pbio.3001922

Simmonds P, Becher P, Bukh J, Gould EA, Meyers G, Monath T, Muerhoff S, Pletnev A, Rico- Hesse R, Smith DB, Stapleton JT, and Ictv Report Consortium. 2017b. ICTV Virus Taxonomy Profile: Flaviviridae. The Journal of general virology 98(1): 2–3. 10.1099/jgv.0.000672

Starr EP, Nuccio EE, Pett-Ridge J, Banfield JF, and Firestone MK. 2019. Metatranscriptomic reconstruction reveals RNA viruses with the potential to shape carbon cycling in soil. Proceedings of the National Academy of Sciences 116(51): 25900–25908. 10.1073/pnas.1908291116

Talbert PB, Armache KJ, and Henikoff S. 2022. Viral histones: pickpocket’s prize or primordial progenitor?. Epigenetics & chromatin 15(1): 21. 10.1186/s13072-022-00454-7

Tamames J, Cobo-Simón M, and Puente-Sánchez F. 2019. Assessing the performance of different approaches for functional and taxonomic annotation of metagenomes. BMC genomics 20(1): 960. 10.1186/s12864-019-6289-6

Ter Horst AM, Nigg JC, Dekker FM, and Falk BW. 2019. Endogenous Viral Elements Are Widespread in Arthropod Genomes and Commonly Give Rise to PIWI-Interacting RNAs. Journal of virology 93(6): e02124–18. 10.1128/JVI.02124-18

Terzian C, Pélisson A, and Bucheton A. 2001. Evolution and phylogeny of insect endogenous retroviruses. BMC evolutionary biology 1: 3. 10.1186/1471-2148-1-3

Tomlinson KR, Pablo-Rodriguez JL, Bunawan H, Nanyiti S, Green P, Miller J, Alicai T, Seal SE, Bailey AM, and Foster GD. 2019. Cassava brown streak virus Ham1 protein hydrolyses mutagenic nucleotides and is a necrosis determinant. Molecular plant pathology 20(8): 1080–1092. 10.1111/mpp.12813

Valverde S, Vidiella B, Montañez R, Fraile A, Sacristán S, and García-Arenal F. 2020. Coexistence of nestedness and modularity in host-pathogen infection networks. Nature ecology & evolution 4(4): 568–577. 10.1038/s41559-020-1130-9

Van den Boogaart KG, and Tolosana-Delgado R. 2008. “compositions”: A unified R package to analyze compositional data. Computers & Geosciences 34(4): 320–338. doi:10.1016/j.cageo.2006.11.017

Veglia AJ, Bistolas KSI, Voolstra CR et al. 2023. Endogenous viral elements reveal associations between a non-retroviral RNA virus and symbiotic dinoflagellate genomes. *Commun*. Biol. 6(1): 566. 10.1038/s42003-023-04917-9

Venables WN, and Ripley BD. 2002. Modern Applied Statistics with S (Fourth edition). New York: Springer. ISBN 0-387-95457-0. Retrieved from https://www.stats.ox.ac.uk/pub/MASS4/

Vieira P, Subbotin SA, Alkharouf N, Eisenback J, and Nemchinov LG. 2022. Expanding the RNA virome of nematodes and other soil-inhabiting organisms. Virus evolution 8(1): veac019. 10.1093/ve/veac019

Wang S, Liu H, Zu X, Liu Y, Chen L, Zhu X, Zhang L, Zhou Z, Xiao G, and Wang W. 2016. The ubiquitin-proteasome system is essential for the productive entry of Japanese encephalitis virus. Virology 498: 116–127. 10.1016/j.virol.2016.08.013

Wasik BR, and Turner PE. 2013. On the biological success of viruses. Annual review of microbiology 67: 519–541. 10.1146/annurev-micro-090110-102833

Weitz JS, Poisot T, Meyer JR, Flores CO, Valverde S, Sullivan MB, and Hochberg ME. 2013. Phage-bacteria infection networks. Trends in microbiology 21(2): 82–91. 10.1016/j.tim.2012.11.003

Wickham H. 2016. ggplot2: Elegant Graphics for Data Analysis. Springer-Verlag New York. ISBN 978-3-319-24277-4, https://ggplot2.tidyverse.org

Wnuk SF, and Robins MJ. 2006. Ribonucleotide reductase inhibitors as anti-herpes agents. Antiviral research 71(2-3): 122–126. 10.1016/j.antiviral.2006.03.002

Wolf YI, Silas S, Wang Y, Wu S, Bocek M, Kazlauskas D, Krupovic M, Fire A, Dolja VV, and Koonin EV. 2020. Doubling of the known set of RNA viruses by metagenomic analysis of an aquatic virome. Nature microbiology 5(10): 1262–1270. 10.1038/s41564-020-0755-4

Wu H, Pang R, Cheng T, Xue L, Zeng H, Lei T, Chen M, Wu S, Ding Y, Zhang J, Shi M, and Wu Q. 2020. Abundant and Diverse RNA Viruses in Insects Revealed by RNA-Seq Analysis: Ecological and Evolutionary Implications. mSystems 5(4): e00039–20. 10.1128/mSystems.00039-20

Wu N, Zhang P, Liu W, and Wang X. 2018. Sogatella furcifera hepe-like virus: First member of a novel Hepeviridae clade identified in an insect. Virus research 250: 81–86. 10.1016/j.virusres.2018.03.018

Wu R, Pisani D, and Donoghue PCJ. 2023. The unbearable uncertainty of panarthropod relationships. Biology Letters 19(1). 10.1098/rsbl.2022.0497

Xia Y, Sun J, Chen DG. 2018. Compositional Analysis of Microbiome Data. In: Statistical Analysis of Microbiome Data with R. ICSA Book Series in Statistics. Springer, Singapore. 10.1007/978-981-13-1534-3_10

Yamasaki H, Fujimoto S, and Miyazaki K. 2015. Phylogenetic position of Loricifera inferred from nearly complete 18S and 28S rRNA gene sequences. Zoological letters 1: 18. 10.1186/s40851-015-0017-0

Zayed AA, Wainaina JM, et al. 2022. Cryptic and abundant marine viruses at the evolutionary origins of Earth’s RNA virome. *Science (New York*, N.Y.) 376(6589): 156–162. 10.1126/science.abm5847

Zhang ZQ. 2013. Animal biodiversity: An outline of higher-level classification and survey of taxonomic richness (Addenda 2013). Zootaxa 3703: 1–82. 10.11646/zootaxa.3703.1.1

Zhang YY, et al. 2022. Viromes in marine ecosystems reveal remarkable invertebrate RNA virus diversity. Sci. China Life Sci 65: 426–437. 10.1007/s11427-020-1936-2

Zhang YZ, Shi M, and Holmes EC. 2018. Using Metagenomics to Characterize an Expanding Virosphere. Cell 172(6): 1168–1172. 10.1016/j.cell.2018.02.043

Zhao S, Guo Y, Sheng Q, and Shyr Y. 2014. Heatmap3: an improved heatmap package with more powerful and convenient features. BMC Bioinformatics 15: P16. 10.1186/1471-2105-15-S10-P16

