## Supplementary material for "Unveiling the Hidden Viromes Across the Animal Tree of Life: Insights from a Taxonomic Classification Pipeline Applied to Invertebrates of 31 Metazoan Phyla": fig_s2.pdf

A

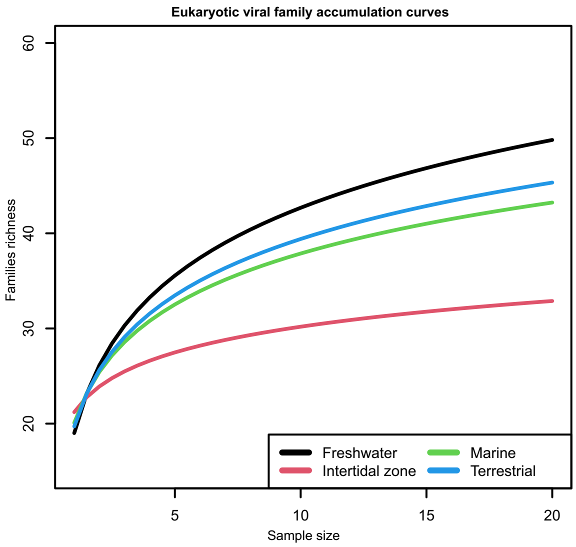


B

| | ANN | ART | BRY | CHO | CNI | CTE | ECH | KIN | MOL | NMD | NME | ONY | PLT | POR | ROT | XEN |
|---|---|---|---|---|---|---|---|---|---|---|---|---|---|---|---|---|
| ANN | 1 | 0.86 | 0.8 | 0.98 | 0.72 | 0.95 | 0.61 | 0.89 | 0.68 | 0.99 | 0.52 | 0.66 | 0.99 | 0.96 | 1 | 0.57 |
| ART | | 1 | 0.93 | 0.99 | 0.99 | 0.96 | 1 | 0.99 | 0.94 | 0.94 | 0.93 | 0.82 | 0.99 | 0.99 | 1 | 0.9 |
| BRY | | | 1 | 0.87 | 0.88 | 1 | 0.86 | 0.46 | 0.32 | 0.77 | 0.74 | 0.94 | 0.98 | 0.66 | 0.97 | 0.78 |
| CHO | | | | 1 | 0.99 | 0.97 | 0.99 | 0.88 | 0.84 | 0.6 | 0.85 | 1 | 0.99 | 0.72 | 0.99 | 0.72 |
| CNI | | | | | 1 | 0.98 | 0.81 | 0.97 | 0.69 | 0.94 | 0.65 | 0.97 | 0.99 | 0.96 | 1 | 0.9 |
| CTE | | | | | | 1 | 0.93 | 0.8 | 0.63 | 0.89 | 0.9 | 0.96 | 1 | 0.74 | 0.99 | 0.81 |
| ECH | | | | | | | 1 | 0.99 | 0.73 | 0.99 | 0.92 | 0.87 | 0.99 | 0.99 | 1 | 0.99 |
| KIN | | | | | | | | 1 | 0.87 | 0.85 | 0.82 | 0.99 | 0.94 | 0.69 | 0.98 | 0.82 |
| MOL | | | | | | | | | 1 | 0.92 | 0.68 | 0.9 | 0.8 | 0.76 | 0.93 | 0.7 |
| NMD | | | | | | | | | | 1 | 0.66 | 0.98 | 0.67 | 0.71 | 0.59 | 0.86 |
| NME | | | | | | | | | | | 1 | 0.83 | 0.95 | 0.99 | 0.99 | 0.72 |
| ONY | | | | | | | | | | | | 1 | 0.99 | 0.99 | 0.99 | 0.83 |
| PLT | | | | | | | | | | | | | 1 | 0.89 | 1 | 0.85 |
| POR | | | | | | | | | | | | | | 1 | 0.9 | 0.93 |
| ROT | | | | | | | | | | | | | | | 1 | 0.98 |
| XEN | | | | | | | | | | | | | | | | 1 |

- Non signficant centroid location difference
- Significant difference in within−group dispersions
- Significant difference in dispersions and possibly in centroid locations
- Significant centroid location difference
