## Supplementary figures and images for "Unveiling the Hidden Viromes Across the Animal Tree of Life: Insights from a Taxonomic Classification Pipeline Applied to Invertebrates of 31 Metazoan Phyla"

### fig_s1.pdf

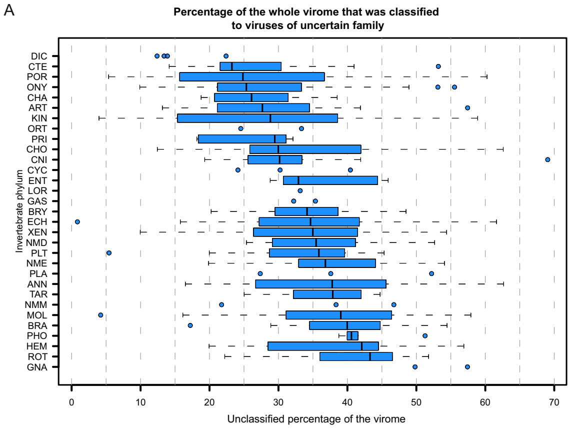

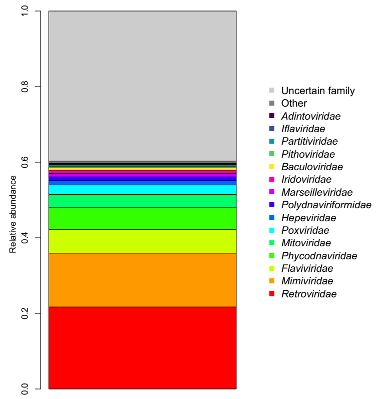
